# Propagation path of a flowering cherry (*Cerasus* × *yedoensis*) cultivar ‘Somei-Yoshino’ traced by somatic mutations

**DOI:** 10.1101/2023.07.11.548633

**Authors:** Kenta Shirasawa, Tomoya Esumi, Akihiro Itai, Katsunori Hatakeyama, Tadashi Takashina, Takuji Yakuwa, Katsuhiko Sumitomo, Takeshi Kurokura, Eigo Fukai, Keiichi Sato, Takehiko Shimada, Katsuhiro Shiratake, Munetaka Hosokawa, Yuki Monden, Makoto Kusaba, Hidetoshi Ikegami, Sachiko Isobe

**Affiliations:** Department of Frontier Research and Development, Kazusa DNA Research Institute, Kisarazu, Japan; Academic Assembly Institute of Agricultural and Life Sciences, Shimane University, Matsue, Japan; Department of Agricultural and Life Science, Kyoto Prefectural University, Kyoto, Japan; Faculty of Agriculture, Iwate University, Morioka, Japan; Horticultural Research Institute, Yamagata Integrated Agricultural Research Center, Sagae, Japan; Yamagata Nishi High School, Yamagata, Japan; Institute of Vegetable and Floriculture Science, NARO, Tsukuba, Japan; Faculty of Agriculture, Utsunomiya University, Utsunomiya, Japan; Graduate School of Science and Technology, Niigata University, Niigata, Japan; Yamanashi Kofu Minami High School, Kofu, Japan; Institute of Fruit Tree and Tea Science, NARO, Shizuoka, Japan; Graduate School of Bioagricultural Sciences, Nagoya University, Nagoya, Japan; Department of Agricultural Science, Kindai University, Nara, Japan; Graduate School of Environmental and Life Science, Okayama University, Okayama, Japan; Graduate School of Integrated Sciences for Life, Hiroshima University, Higashi-Hiroshima, Japan; Resident, Fukuoka, Japan

**Keywords:** Clone, flowering cherry, genome sequence, somatic mutation, Somei-Yoshino

## Abstract

Flowering cherry cultivar ‘Somei-Yoshino’ (*Cerasus* × *yedoensis*) has been clonally propagated and spread all around the world including Japan. ‘Somei-Yoshino’ is thought to be an interspecific hybrid derived from *C. spachiana* and *C. speciosa*; however, its origin is unclear. Since somatic mutations are randomly induced in genomes and stably transmitted through generations, we aimed to identify somatic mutations in the genome of ‘Somei-Yoshino’ to trace its propagation path. A total of 46 ‘Somei-Yoshino’ clones were collected from all over Japan and subjected to whole-genome sequencing. The results revealed 684 single nucleotide variants, of which 71 were found in more than two clones. Clustering analysis of the 46 clones using these 71 variants revealed six groups, four of which contained 40 of the 46 clones. In addition, because each of the four clones closely planted in Ueno Park, Tokyo, Japan, clustered into the four different groups, we considered that these four clones could be the ancestors of the ‘Somei-Yoshino’ clones found in Japan. Furthermore, based on the comparison of mutant alleles with the genomes of *Cerasus* species, one of the four trees was concluded as the closest to the origin. Here, we propose that the origin of ‘Somei-Yoshino’ is a chimera derived from at least four somatic mutants.

## Introduction

‘Somei-Yoshino’ (*Cerasus* × *yedoensis*) is one of the most popular flowering cherry cultivars distributed all over the world. The flowers of ‘Somei-Yoshino’ are beautiful and are considered a symbol of spring by most. Therefore, methods have been developed to forecast flowering cherry bloom based on temperature or gene expression (Aono and Murakami 2017; Shirasawa et al. 2022). ‘Somei-Yoshino’ is self-incompatible and has a highly heterozygous genome, owing to its origin as an interspecific hybrid of *Cerasus spachiana* and *Cerasus speciosa* (Takenaka 1963; Innan et al. 1995; Shirasawa et al. 2019), and thus cannot be propagated by seeds. Therefore, ‘Somei-Yoshino’ has been clonally propagated by grafting or cutting methods (Iketani et al. 2007). Nevertheless, whether ‘Somei-Yoshino’ is a natural or an artificial hybrid remains unknown except for a tale that ‘Somei-Yoshino’ trees were first sold by a gardener(s) at Somei village (Tokyo, Japan) at the end of the Edo period. Some studies suggest that the original predecessor tree of ‘Somei-Yoshino’ was planted in Koishikawa Botanical Garden (Tokyo, Japan) (Nakai 1935), Ueno Park (Tokyo, Japan) (Nakamura et al. 2015), or Kaiseizan Park (Fukushima, Japan) (Sampei 2018). A recent study suggested that four trees (#133, #134, #136, and #138) closely planted in Ueno Park could be candidates for the original predecessor tree of ‘Somei-Yoshino’ (Nakamura I, Tsuchiya A, Takahashi H, Makabe S 2015). Therefore, the origin of ‘Somei-Yoshino’ remains of interest to researchers as well as to people fascinated with cherry blossoms.

Somatic mutations occur randomly in different organs of an individual and cannot be reversed, which leads to chimerism (Wang et al. 2019; Orr et al. 2020; Hofmeister et al. 2020). Some of the mutations in gene bodies affect gene functions, leading to phenotypic alterations (Kobayashi et al. 2004). In trees, bud sports are the result of somatic mutations. Since bud sports are genetically stable, grafting or cutting is used to propagate the mutants as new cultivars (Sanada et al. 1993). Furthermore, silent somatic mutations, which do not affect gene function nor cause phenotypic variations, also exist in genomes and are stably inherited. Therefore, somatic mutations could be used to trace the propagation path of clonally propagated organisms.

To the best of our knowledge, the clonality of ‘Somei-Yoshino’ has been investigated via DNA fingerprinting using microsatellite markers (Innan et al. 1995; Iketani et al. 2007). In addition, somatic mutations within a single ‘Somei-Yoshino’ tree have been studied using techniques such as temperature gradient gel electrophoresis (TGGE) (Diwan et al. 2014) and double-digest restriction site-associated DNA sequencing (ddRAD-Seq) (Ueno et al. 2023). These studies indicated that the genetic identity of ‘Somei-Yoshino’ clones is high, although a few somatic mutations could be detected. Since the genome of ‘Somei-Yoshino’ has been sequenced (Shirasawa et al. 2019), a large-scale mutation analysis could be conducted via whole-genome resequencing to identify genome-wide somatic mutations and to trace the propagation path of ‘Somei-Yoshino’. Here, we analyzed the whole-genome sequence of 46 ‘Somei-Yoshino’ clones to detect somatic mutations.

The 46 clones were clustered into six groups, based on nucleotide sequence variants. This result suggests that the ‘Somei-Yoshino’ clones originated from four genotypes planted within a small area of Ueno Park (Tokyo, Japan).

## Materials and methods

### Plant material

Leaves were collected from 46 randomly selected ‘Somei-Yoshino’ trees grown in 19 prefectures of Japan (Table S1, Fig. S1); these 46 trees included the four trees, #133, #134, #136, and #138, planted in Ueno Park (Tokyo, Japan). Genomic DNA was extracted from the leaves using the FavorPrep Plant Genomic DNA Extraction Mini Kit (Favorgen, Ping-Tung, Taiwan).

### DNA sequencing

Genomic DNA libraries were prepared with a PCR-free method using the Swift 2S Turbo Flexible DNA Library Kit (Swift Biosciences, Ann Arbor, MI, USA), and converted into a DNA nanoball sequencing library with the MGI Easy Universal Library Conversion Kit (MGI Tech, Shenzhen, China). The library was sequenced on the DNBSEQ G400RS (MGI Tech) instrument in paired-end, 150 bp mode. The sequence data of tree #136, with accession numbers DRR169775 (SyTKY0) and DRR169776 (SyTKY1), were obtained from a DNA database.

### Detection and analysis of somatic mutations

Low-quality bases (quality score < 10) and adaptor sequences (AGATCGGAAGAGC) were trimmed with PRINSEQ (Schmieder and Edwards 2011) and fastx_clipper, respectively, in FASTX-Toolkit (http://hannonlab.cshl.edu/fastx_toolkit), and the remaining high-quality reads were mapped on to the haplotype-resolved genome sequence of ‘Somei-Yoshino’ (CYE_r3.1.pseudomolecule) with Bowtie 2 (Langmead and Salzberg 2012). The resultant SAM files were converted into BAM files with the *view* command of SAMtools (Li 2011). The gVCF files were generated from the BAM files using the *mpileup* command (-Ou -a DP,AD,INFO/AD) and *call* command (-Oz -m -g 0,10) in BCFtools (version 1.9) (Li 2011), and normalized using the *norm* command of BCFtools. The individual gVCF files were merged into a single VCF file using the *merge* command of BCFtools. High-confidence, biallelic, homozygous single nucleotide variants (SNVs) were selected with VCFtools (Danecek et al. 2011). Copy number variations (CNVs) were detected with cnv-seq (Xie and Tammi 2009), and the effect of sequence variations on gene function was predicted with SNPeff (Cingolani et al. 2012).

To identify cultivars, the ddRAD-Seq data of 139 lines were downloaded from a DNA database (GenBank accession numbers: DRR169804–DRR169942). High-quality reads, selected as described above, were mapped on to the genome sequence of ‘Somei-Yoshino’ (CYE_r3.1.pseudomolecule), and SNVs were detected as described previously (Shirasawa et al. 2019). SNVs identified in the 46 samples, based on the analysis of ddRAD-Seq reads and gVCF files, were combined to calculate the distance between each pair of 185 samples (= 139 + 46), and a neighbor-joining tree was created with Tassel 5 (Bradbury et al. 2007).

To identify the possible ancestral alleles of SNVs, whole-genome sequence reads obtained from 10 lines belonging to six *Cerasus* species (*C. campanulata, C. serrulata, C. spachiana, C. speciosa, C. jamasakura*, and *C*. × *nudiflora*) were obtained from a public DNA database (GenBank accession numbers: DRR169795–DRR169803 and SRR6957274). SNV detection was performed as above, and the SNV genotypes identified in the 56 samples were compared.

## Results

### Cultivar identification

Whole-genome sequence reads obtained from 46 samples in this study (Table S1, Fig. S1) and ddRAD-Seq reads obtained from 139 lines in our previous study, including ‘Somei-Yoshino’ (Cerasus_1-72), were aligned on to the reference genome sequence of ‘Somei-Yoshino’. A total of 5,328 SNPs were detected across the 185 samples, and genetic distances between each pair of the 185 samples were calculated to create a dendrogram (Fig. 1a). As expected, the 46 samples and ‘Somei-Yoshino’ (Cerasus_1-72) clustered together, with an average genetic distance of 0.01, while the other lines including probable sister and parent lines formed a separate cluster, with a genetic distance of 0.37. We concluded that the 46 samples from this study were indeed ‘Somei-Yoshino’ clones.

**Fig. 1.**
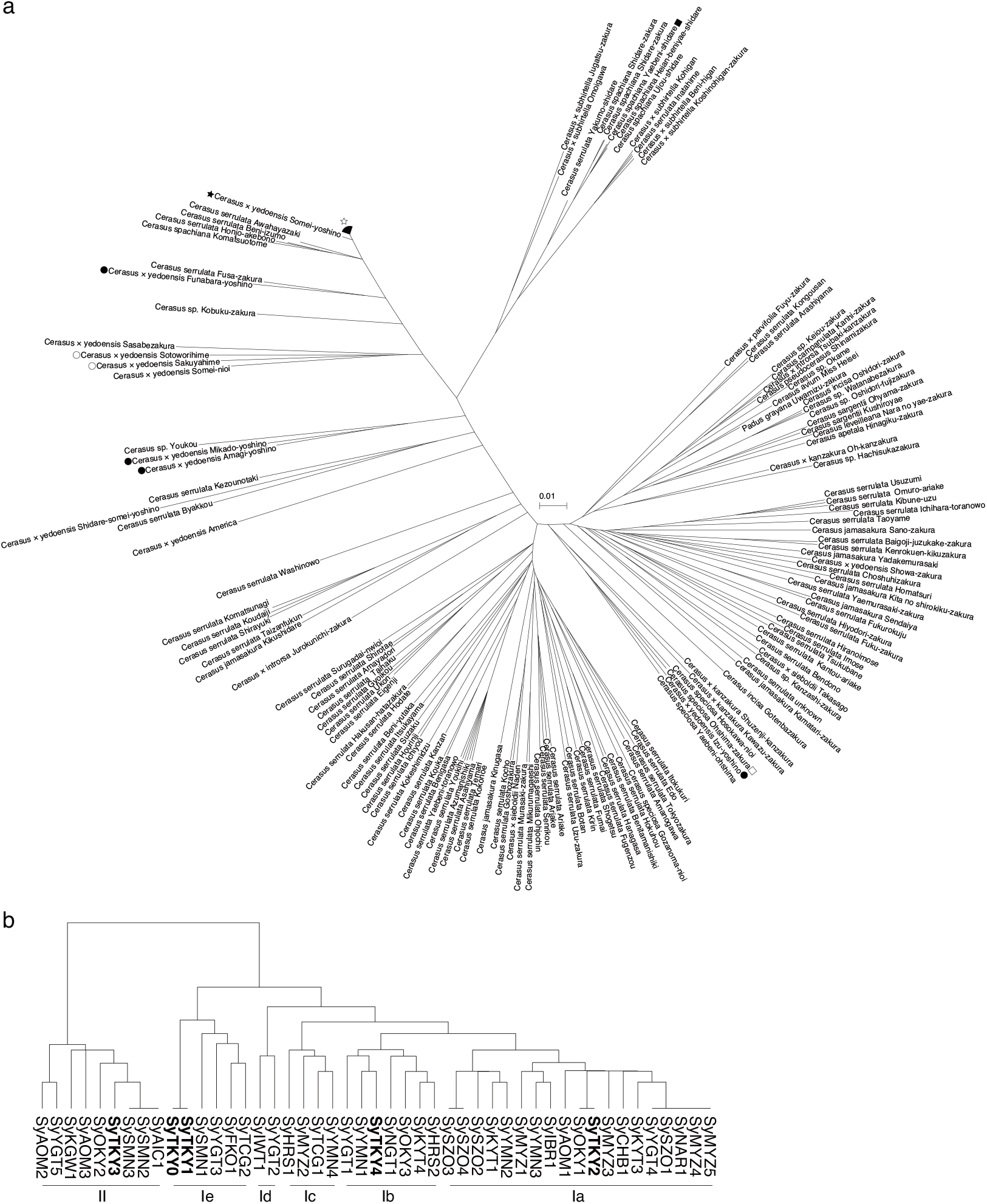
Dendrograms of flowering cherry.**a** Dendrogram of 46 ‘Somei-Yoshino’ clones and 139 cherry lines. Black and white stars indicate the ‘Somei-Yoshino’ tree (Cerasus 72-1) used in our previous study (Shirasawa et al. 2019) and 46 clones used in this study, respectively. Black circles show synthetic hybrids generated by crossing *C. spachiana* and *C. speciosa* (Takenaka 1963). Black and white squares indicate probable parental lines of *C. spachiana* and *C. speciosa*, respectively. **b** Dendrogram of 46 ‘Somei-Yoshino’ clones. Boldface indicate four trees planted in Ueno Park (Tokyo, Japan). SyTKY0 and SyTKY1 represent different sequence datasets obtained from the same tree (#136). Group names (Ia–Ie and II) are shown below the dendrogram.

### Detection and characterization of somatic mutations

Based on the whole-genome sequence analysis, 80,334 sequence variant candidates were detected in the 46 ‘Somei-Yoshino’ samples. First, we selected 35,757 biallelic SNVs. Next, 1,942 homozygous sites were selected because the two haplotype sequences of the ‘Somei-Yoshino’ genome have been resolved, on which the heterozygous variants were not allowed, probably due to errors that occurred in alignments or sequencing. Then, 1,749 single nucleotide variants, whose genotypes were consistent among the biological replicates of SyTKY0 and SyTKY1, were retained. Finally, we applied two filtering criteria, namely, read depth (≤50) and quality (≥80), to select 684 high-confidence SNVs, which were evenly distributed across the genome (Fig. 2). The number of variants across the 46 samples was 50.3, on average, with the maximum value of 144 in SyAOM1, followed by 106 in SyYMN3 and 90 in SyKGW1 (Fig. 3). The 684 SNVs consisted of 285 C/G to T/A transitions (41.7%), 122 A/T to T/A transversions (17.8%), 120 A/T to G/C transitions (17.5%), 72 C/G to A/T transversions (10.5%), 51 C/G to G/C transversions (7.5%), and 34 A/T to C/G transversions (5.0%). The transition/transversion ratio was 1.45. Among the 684 variants, 88 variants (12.9%) were located in gene bodies, whereas the remaining 596 variants (87.1%) were found in intergenic regions.

**Fig. 2.**
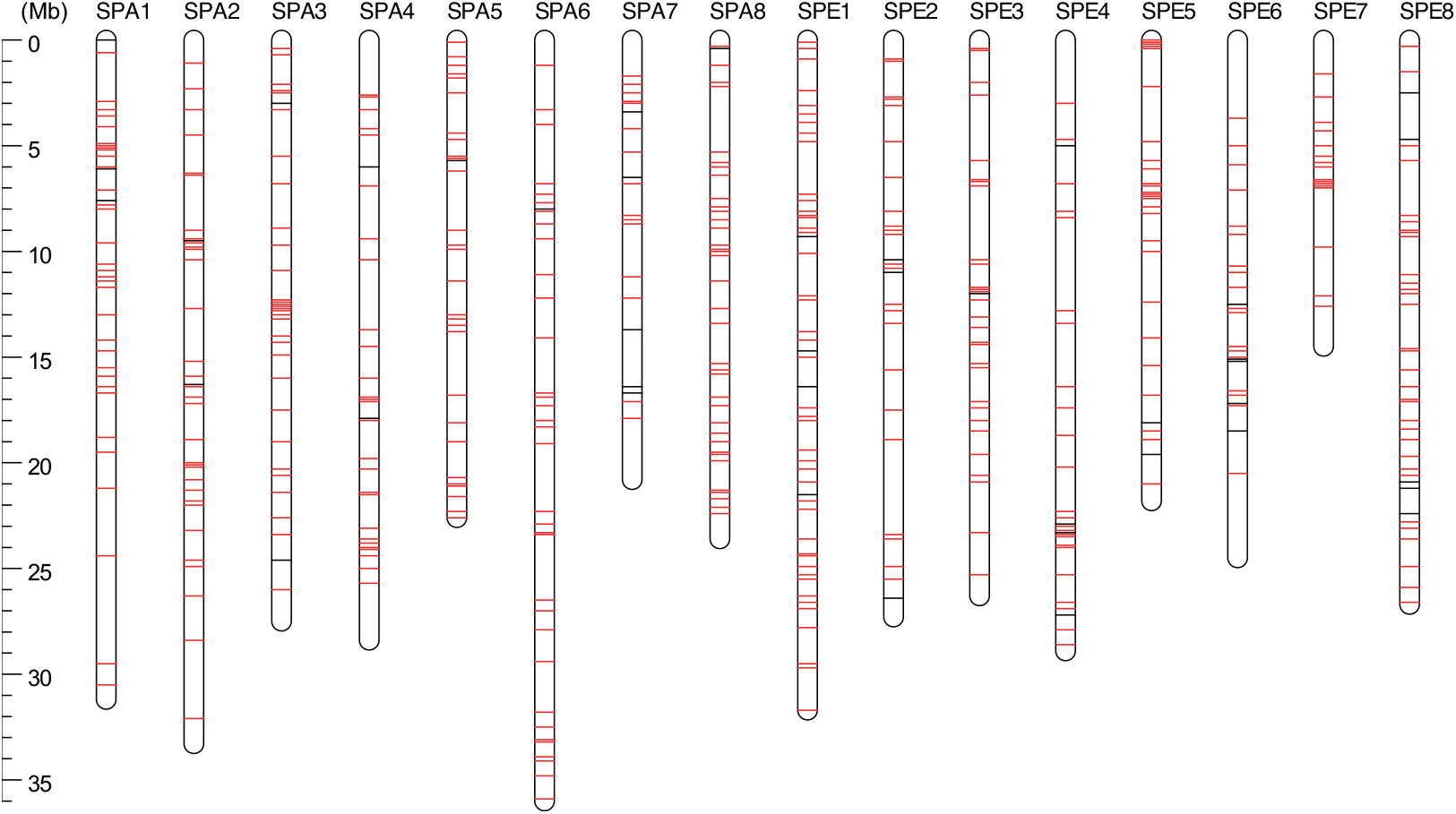
Genomic position of somatic mutations. SPA1–SPA8 and SPE1–SPE8 indicate the chromosomes of ‘Somei-Yoshino’. Black and red lines on the chromosome show common and unique variants, respectively, across the 46 clones.

**Fig. 3.**
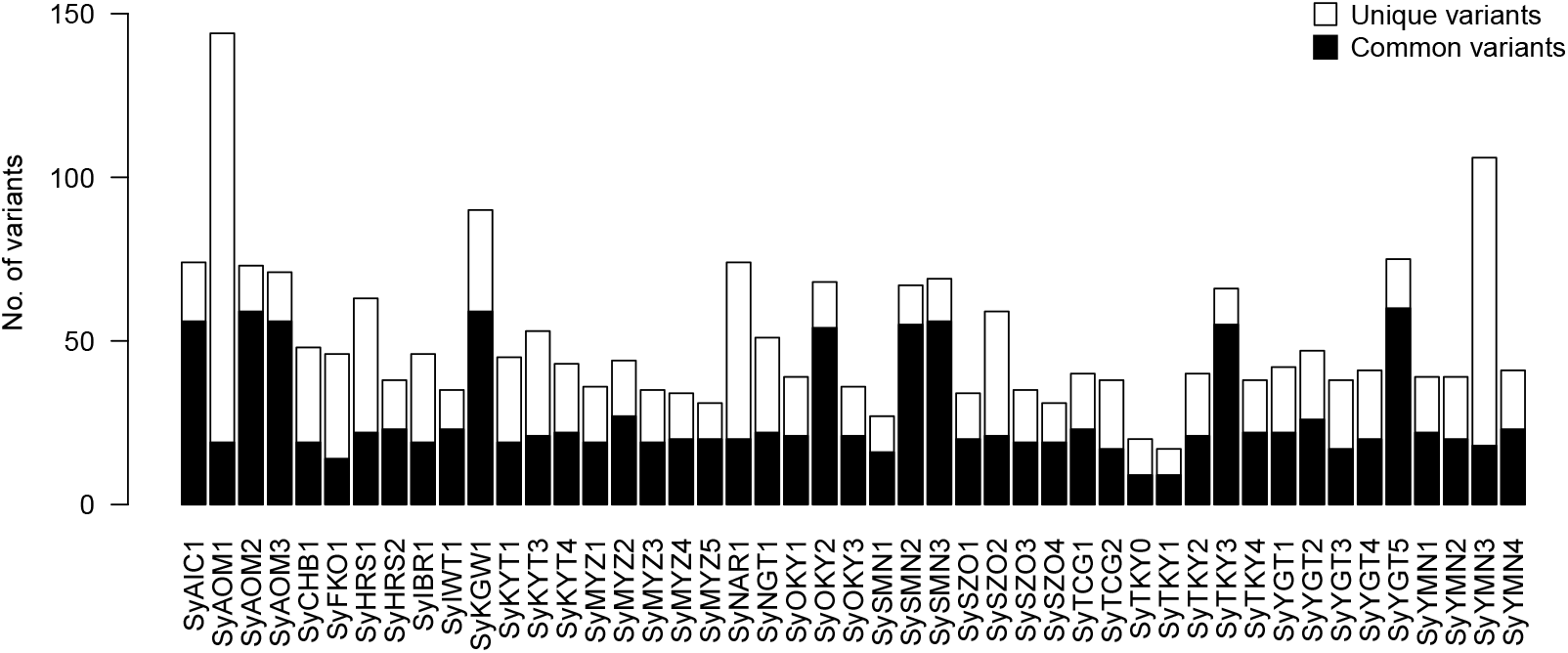
Numbers of variants in the 46 ‘Somei-Yoshino’ clones. Black and white bars show numbers of common and unique variants, respectively.

Of the 684 variants, 613 were unique to a single clone, and 71 were common to at least two clones (Fig. 3). Of the 88 variants identified in gene bodies, seven variants, all of which were unique to a single clone, were predicted to have a high impact on gene functions (Table 1, Fig. S2). These seven variants included four non-sense mutations (in CYE_r3.1SPA5_g009120 in SyYMN2, CYE_r3.1SPE0_g067760 in SyYGT3, CYE_r3.1SPE5_g007060 in SyIBR1, and CYE_r3.1SPE8_g002010 in SyYMN3), two mutations at splice acceptor sites (in CYE_r3.1SPA4_g020770 in SyNGT1 and CYE_r3.1SPA5_g023100 in SyYGT4), and one at a splice donor site (in CYE_r3.1SPE0_g037740 in SyYMN3). Moderate-impact variants, all of which were missense mutations, were found in 23 genes (Fig. S2).

**Table 1.**
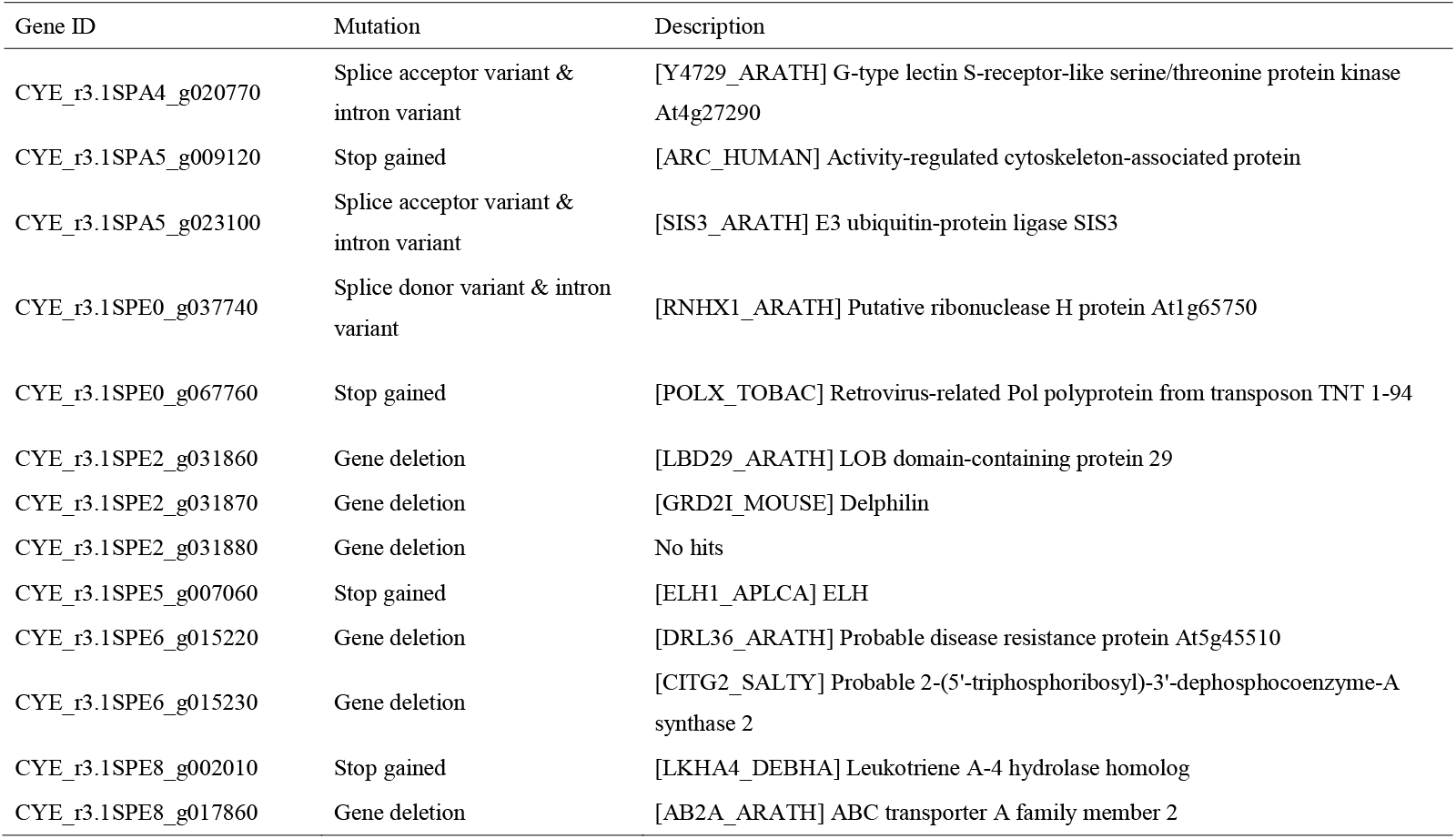
Somatic mutations highly affecting gene functions

### CNVs among the ‘Somei-Yoshino’ clones

At least five CNVs were found in four clones, SyAOM1, SyFKO1, SyYGT2, and SyYMN3 (Fig. 4, Table 1). All five CNVs were deletion mutations with respect to SyTKY1 as a standard; SyAOM1 carried two CNVs, one on chromosome SPE2 (∼25 kb deletion) and another on chromosome SPE8 (∼10 kb deletion), while each of the three remaining clones, SyFKO1, SyYGT2, and SyYMN3, carried a single deletion on chromosomes SPA2 (∼15 kb deletion), SPE6 (∼10 kb deletion), and SPE2 (∼25 kb deletion), respectively. Among these CNVs, the deletions on chromosome SPE2 in SyAOM1 and SyYMN3 might be identical. The deletions encompassed three genes on chromosome SPE2 (CYE_r3.1SPE2_g031860, CYE_r3.1SPE2_g031870, and CYE_r3.1SPE2_g031880), one gene on SPE8 (CYE_r3.1SPE8_g017860), and two genes on SPE6 (CYE_r3.1SPE6_g015220 and CYE_r3.1SPE6_g015230), while the deletion on SPA2 contained no gene.

**Fig. 4.**
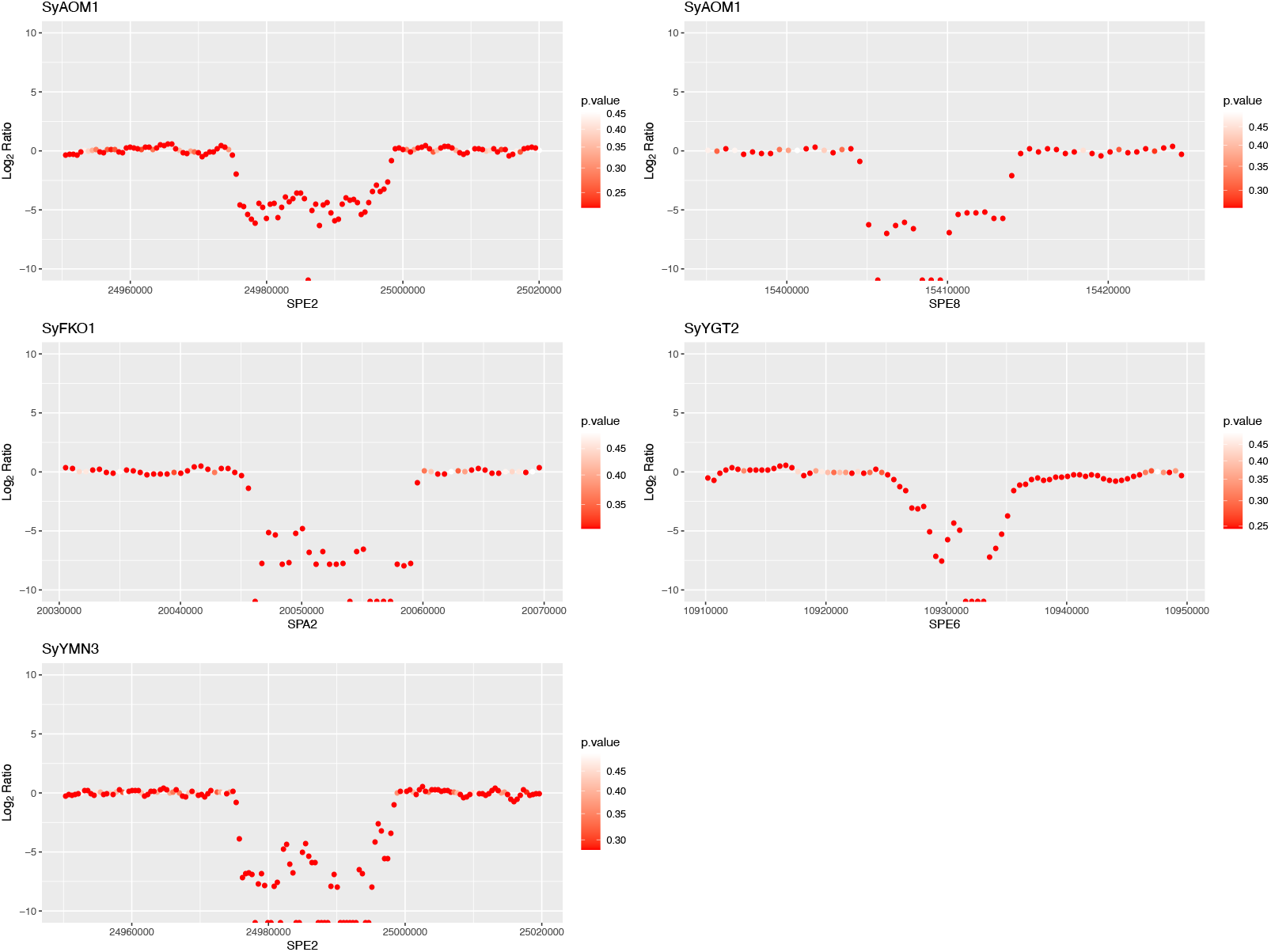
Copy number variations (CNVs) detected in the four ‘Somei-Yoshino’ clones collected from Ueno Park. Clone and chromosome names are shown above and below the plots, respectively.

### Clustering analysis of ‘Somei-Yoshino’ clones

Based on 71 common variants, the 46 samples were clustered into two major groups (I and II), and group I was further divided into five subgroups (Ia to Ie) (Fig. 1b). The numbers of clones in each group were as follows: 19 (Ia), 7 (Ib), 4 (Ic), 2 (Id), 5 (Ie), and 9 (II). These clusters showed no correlation with the sample collection site. Each of the four clones collected from Ueno Park (Tokyo, Japan), SyTKY1–SyTKY4, grouped into four different clusters (Ia, Ib, Ie, and II), which contained 40 of the 46 clones tested.

To identify the group close to the potential origin of ‘Somei-Yoshino’, alleles of the 71 common variants in 46 samples were compared with those of 10 *Cerasus* lines belonging to six species. Since no variations were observed at the 71 sites of the 10 lines, the alleles possessed by the 10 lines were presumed to be the ancestral alleles of *Cerasus*. The median values of the ancestral allele frequency in the six groups were 0.915 (Ia), 0.873 (Ib), 0.866 (Ic), 0.830 (Id), 0.859 (Ie), and 0.408 (II) (Fig. 5a). At the sample level, the ancestral allele frequency ranged from 0.449 (SyYGT5) to 0.969 (SyYMN3) (Fig. 5b).

**Fig. 5.**
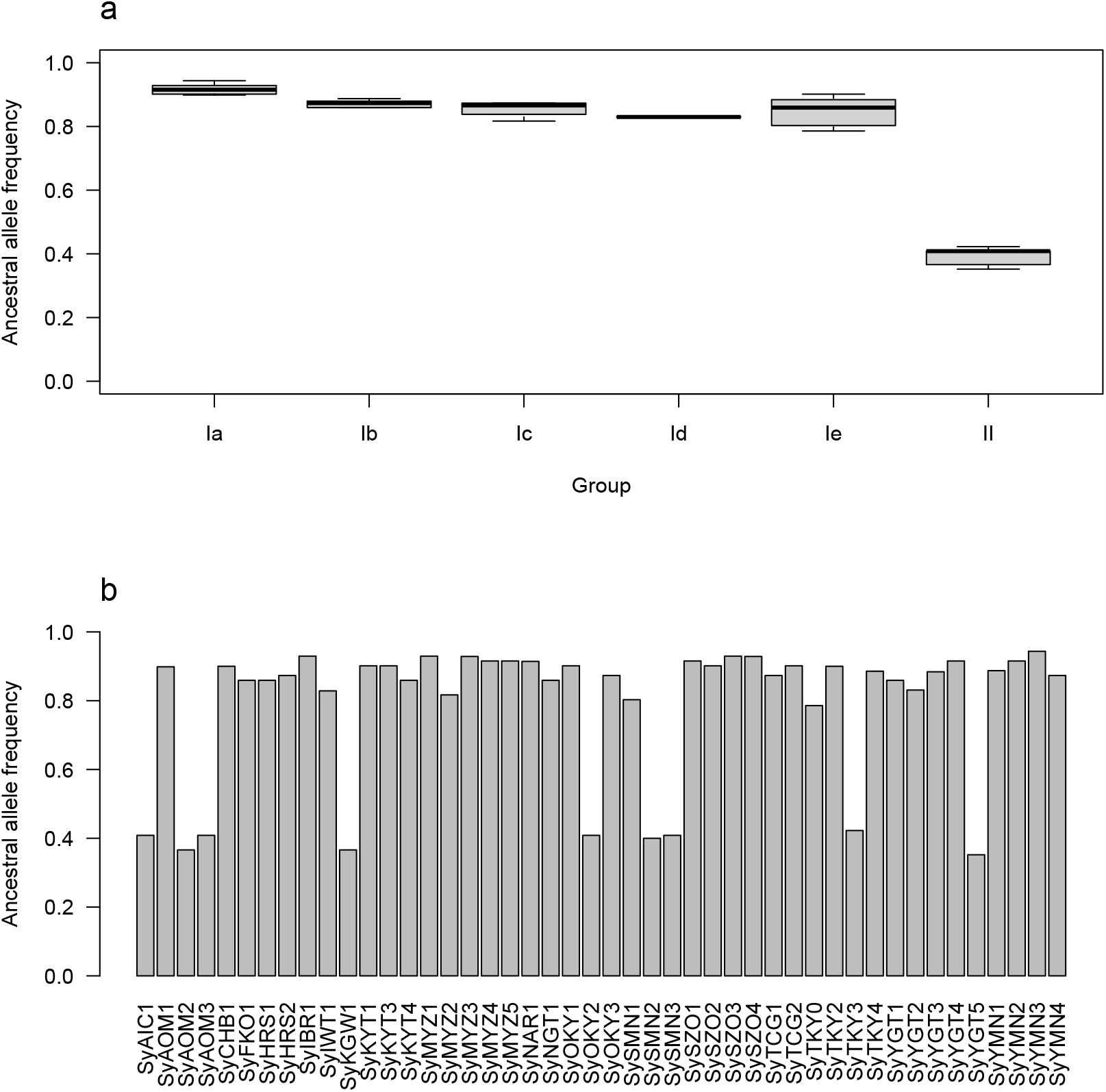
Ancestral allele frequency ‘Somei-Yoshino’ clones. **a** Ancestral allele frequency in the six groups of ‘Somei-Yoshino’ clones. **b** Ancestral allele frequency of ‘Somei-Yoshino’ clones at the sample level.

## Discussion

A total of 684 somatic mutations were detected across 46 ‘Somei-Yoshino’ trees (Fig. 2, Fig. 3). Because this mutation frequency was quite lower than that found among the flowering cherry accessions, the 46 trees were identified as clones of ‘Somei-Yoshino’ (Fig. 1a). Of the 684 mutations, 71 were shared by multiple clones (Fig. 3). Since somatic mutations are not reversible, each of these 71 mutations would have a common origin and would be distributed across multiple lines via clonal propagation. Based on this observation, the 46 ‘Somei-Yoshino’ clones were clustered into six groups, Ia–Ie and II (Fig. 1b). No correlation was detected between the groups and the geographical location of the trees, suggesting that the ‘Somei-Yoshino’ trees were artificially, not naturally, distributed across Japan. On the other hand, interestingly, four trees collected from Ueno Park were classified into four different groups (Fig. 1b). Furthermore, 36 clones in addition to the four trees were included in the four groups (Fig. 1b). The ‘Somei-Yoshino’ clones tested in this study were randomly collected from different locations across Japan, which might imply that most of the ‘Somei-Yoshino’ trees in Japan cluster into four groups. The four trees collected from Ueno Park could represent the origin of ‘Somei-Yoshino’. Among the four trees, SyTKY2 (tree ID #133), which possessed the highest number of ancestral alleles (Fig. 5), was presumed to be the closest to the origin. Although the actual origin was still unclear, we obtained a key set of somatic mutations to identify the origin. We hypothesize that the original tree was a chimera composed of the somatic mutations found in the four groups. Since there are many candidates for the origin of ‘Somei-Yoshino’ in Japan (Iwasaki 1989) as well as in Koishikawa Botanical Garden (Tokyo, Japan) (Nakai 1935) and Kaiseizan Park (Fukushima, Japan) (Sampei 2018), the origin could be discovered by finding the chimera.

The detected mutations consisted of 405 transitions and 279 transversions, with the transition/transversion ratio of 1.45. Among the different mutation types, the C/G to T/A transitions were the most prominent, the proportion of which (41.7%) was comparable with that of somatic mutations in popular (Hofmeister et al. 2020) and ethyl methanesulfonate (EMS)-induced artificial mutations in tomato (Shirasawa et al. 2016). In addition, large-scale deletions (10–25 kb) were also found as somatic mutations, which have been found in not only chemical mutagenesis but also physical mutagenesis studies (Shirasawa et al. 2016; Ichida et al. 2019). Owing to these mutations, the functions of at least 13 genes were severely lost in the clones (Table 1). These mutations could provide phenotypic diversity even in clonally propagated ‘Somei-Yoshino’. In several vegetatively propagated crops, mainly fruit trees, bud sports caused by somatic mutations were reported and used as new cultivars (Sanada et al. 1993). For example, in grape, a transposable element was reported as an inducer of a bud sport, in which the berry skin color was changed from black to white (Kobayashi et al. 2004). While few reports on phenotypic variations are available in ‘Somei-Yoshino’, we observed that the flowering date of #133 in Ueno Park (Tokyo, Japan) was later than those of #134, #136, and #138 in 2023 (data not shown). Further investigation would be required to clarify whether the phenotypic variations are caused by genetic factors (somatic mutation) and/or environmental conditions.

The number of somatic mutations in the 46 trees ranged from 20 (in SyTKY0) to 144 (in SyAOM1) (Fig. 3). This variation in the number of somatic mutations was likely reflected by the number of unique variants rather than that of common variants, even though common variants were more frequent in group II than in group I (Fig. 3). The number of common variants might indicate the time of divergence from the ancestor. In this study, it could be assumed that the branch corresponding to group II is older than those corresponding to the other groups. On the other hand, the number of unique variants might indicate the age of the clone after its propagation via cutting or grafting. It is believed that SyAOM1 is the oldest ‘Somei-Yoshino’ clone planted in 1888 (Iwasaki 1989). Although the SyAOM2 tree is thought to be as old as the SyAOM1 trees, the number of unique variants in SyAOM2 was much less than that in SyAOM1 (Fig. 3). SySZO1 tree is believed to have been obtained from Washington DC, USA, where ‘Somei-Yoshino’ trees from Japan were planted in 1912, but the number of unique variants in SySZO1 was similar to that in other clones. In addition, although SyHRS2 is thought to have survived the atomic bomb attack in 1945 during World War II, the number of somatic mutations in this clone is not high. Overall, the relationship between the number of mutations and the age of ‘Somei-Yoshino’ was unclear.

In summary, this study demonstrates that the propagation path of clonally propagated plants could be traced by identifying the somatic mutations in the plant genome. This somatic mutation-based tracing method could be used to ensure the quality control of vegetatively propagated crops such as orange, apple, grape, strawberry, sweetpotato, and tea, while protecting the rights of breeders, since these crops are frequently transported both within and across regions.

## Acknowledgments

We thank Prof. O. Arakawa (Hirosaki University, Japan) and two anonymous contributors (A.T. and M.T.) for collecting leaf samples; Hirosaki Park (Aomori, Japan), Takamatsu Park (Iwate, Japan), Kajo Park (Yamagata, Japan), Ueno Park (Tokyo, Japan), Kokubo Park (Yamanashi, Japan), Kumano Shrine (Yamagata, Japan), Okayama Prefectural Multipurpose Grounds (Okayama, Japan), Toryo Park (Kagawa, Japan), Maiduru Park (Fukuoka, Japan), and Mochio Park (Miyazaki, Japan) for providing leaf samples; Prof. K. Okada (Ryukoku University, Japan) and Dr. T. Yukawa (National Museum of Nature and Science, Japan) for insightful discussions; and Y. Kishida, C. Minami, K. Ozawa, H. Tsuruoka, and A. Watanabe (Kazusa DNA Research Institute) for technical assistance.

## Accession numbers

The sequence reads are available from the DDBJ Sequence Read Archive (DRA) under the accession numbers DRR493485–DRR493529 of BioProject PRJDB16216.

