## Supplementary Figure for "Propagation path of a flowering cherry (*Cerasus* × *yedoensis*) cultivar ‘Somei-Yoshino’ traced by somatic mutations"

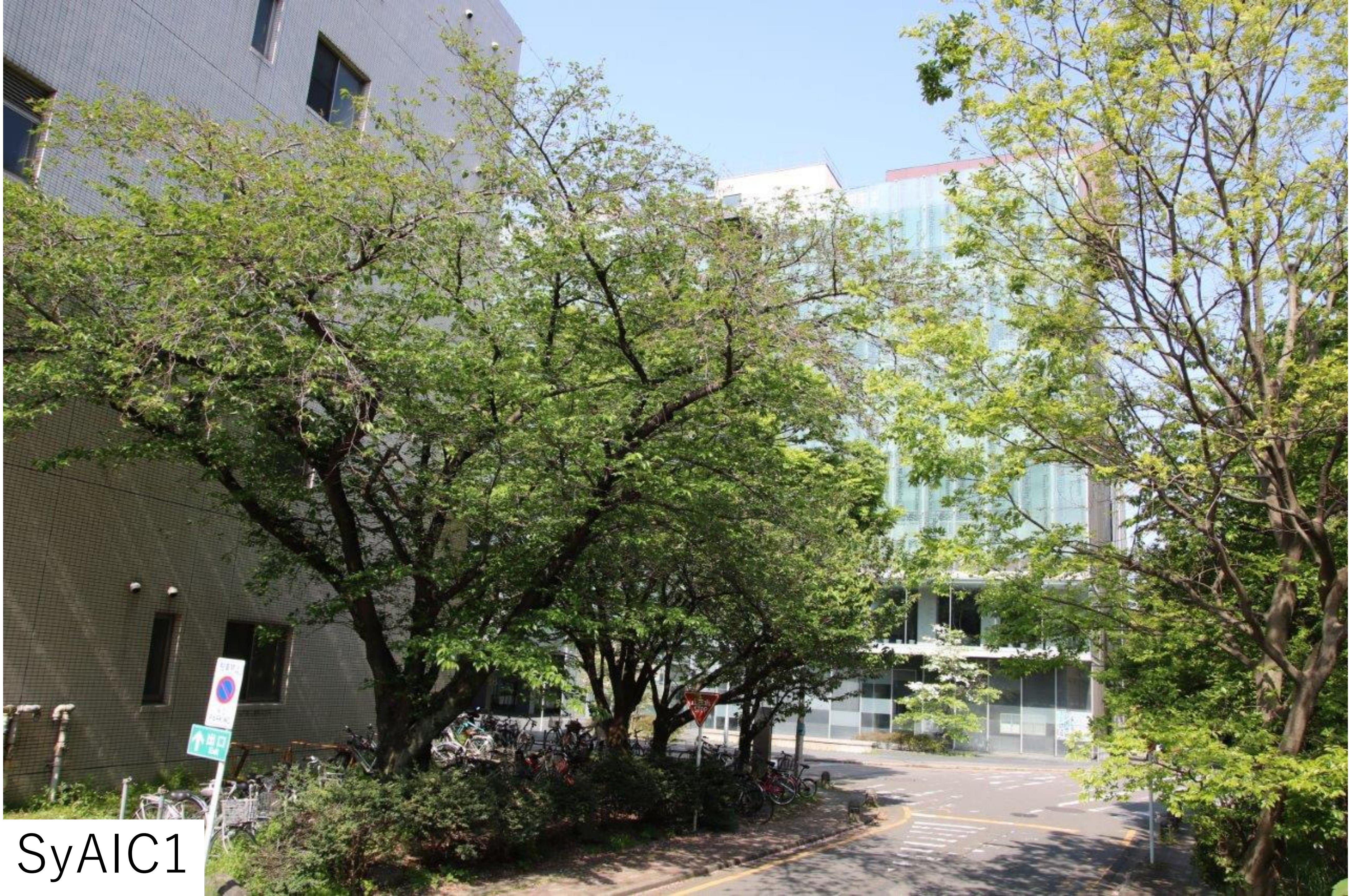

SyAIC1

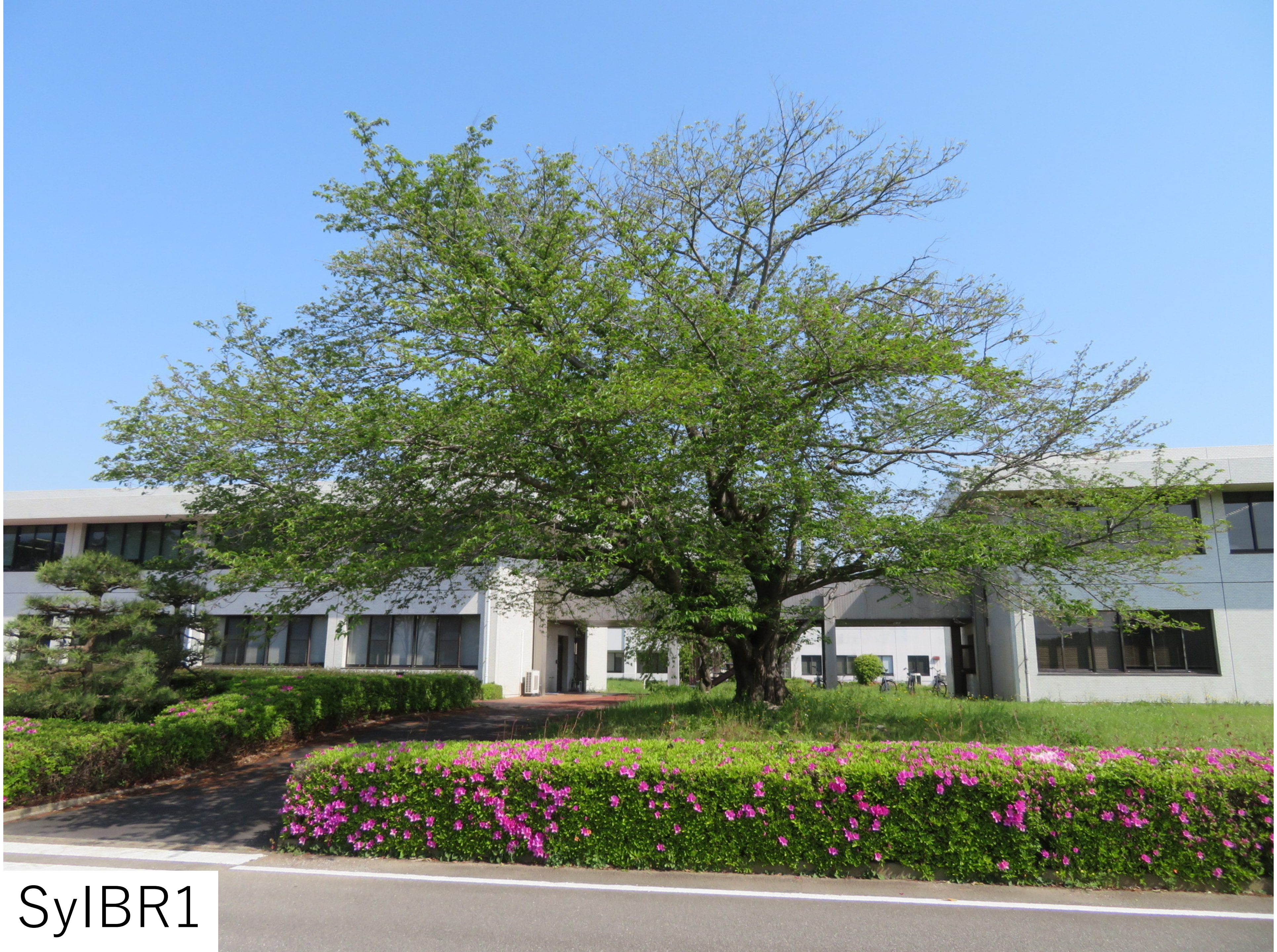

SyIBR1

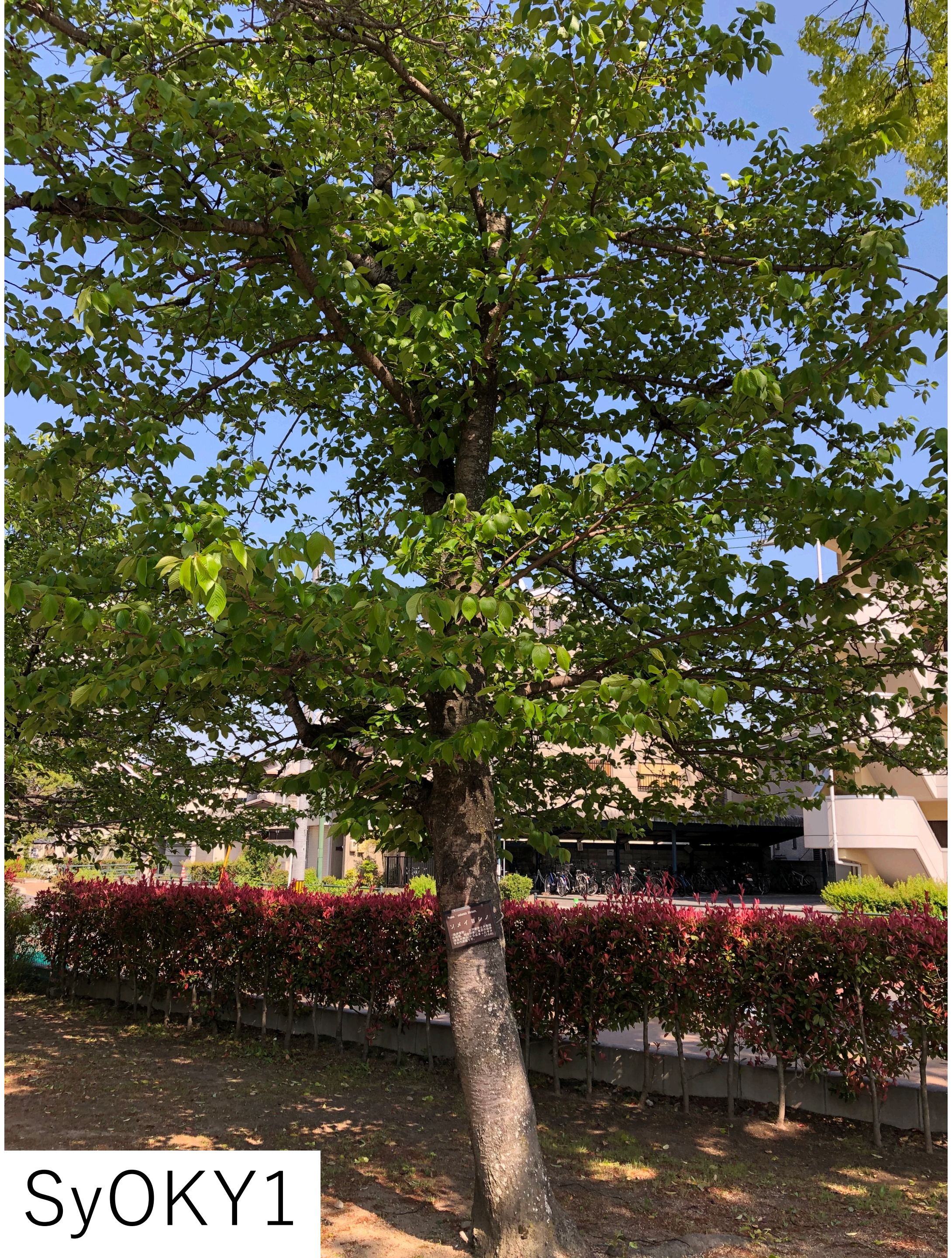

SyOKY1

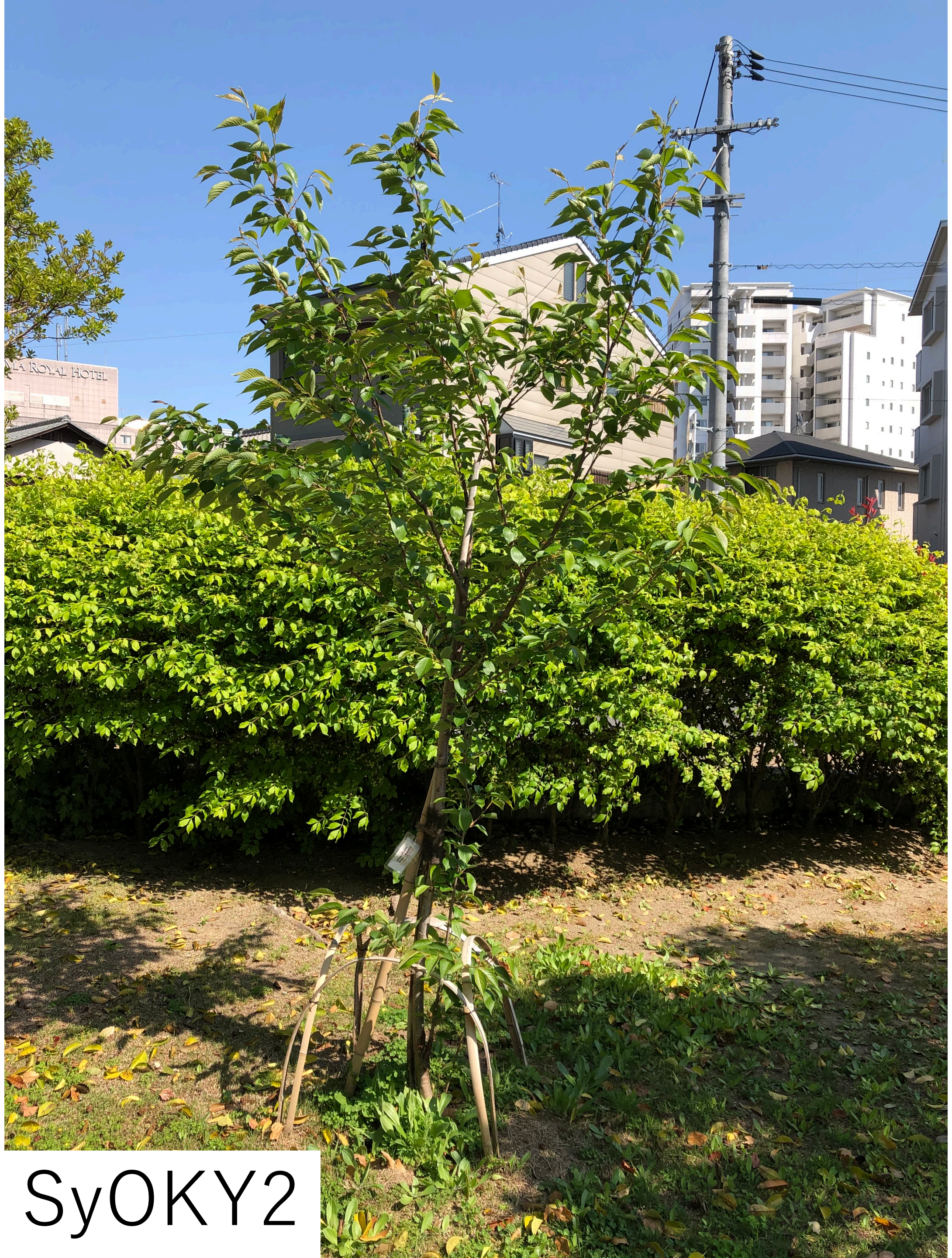


SyOKY2

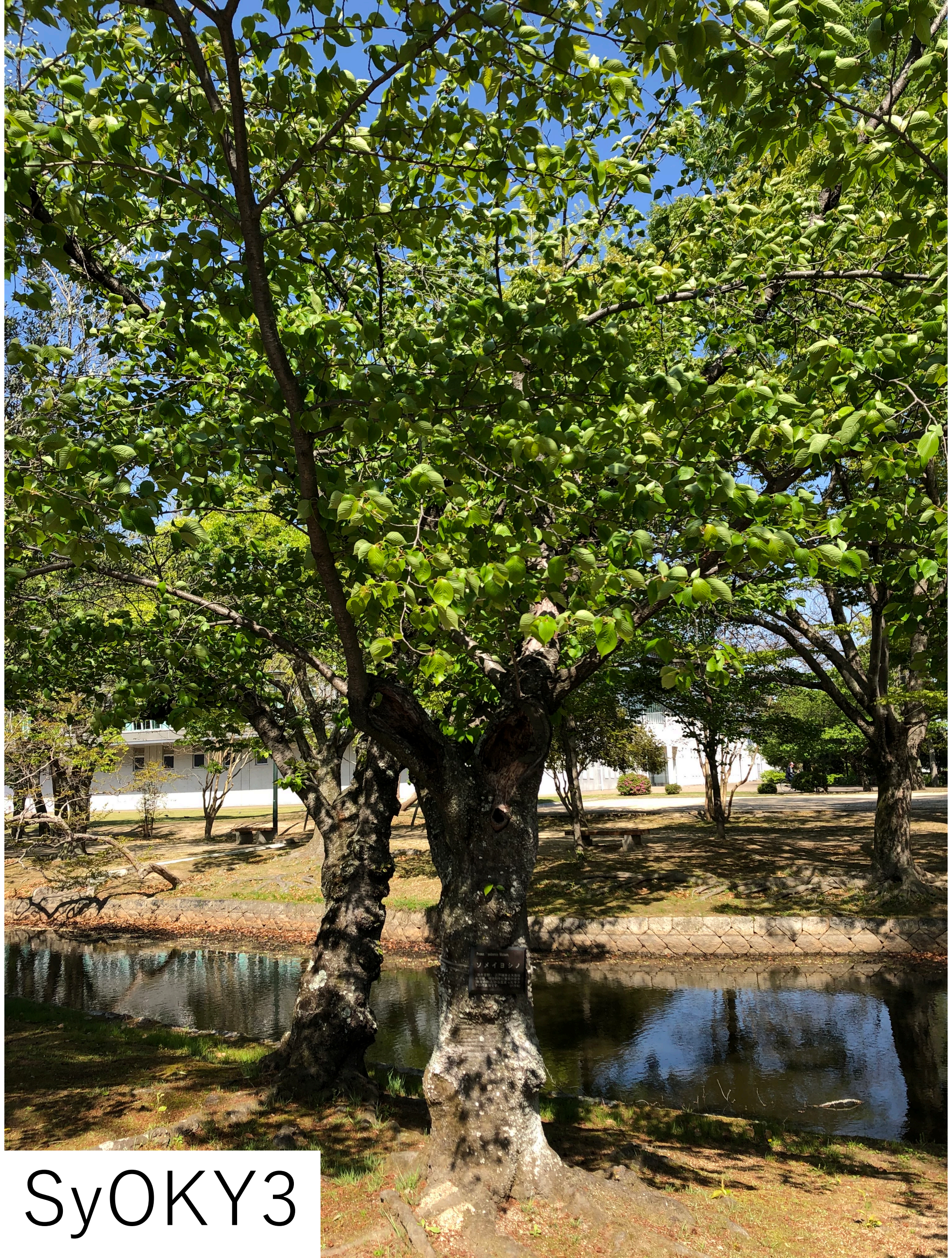

SyOKY3

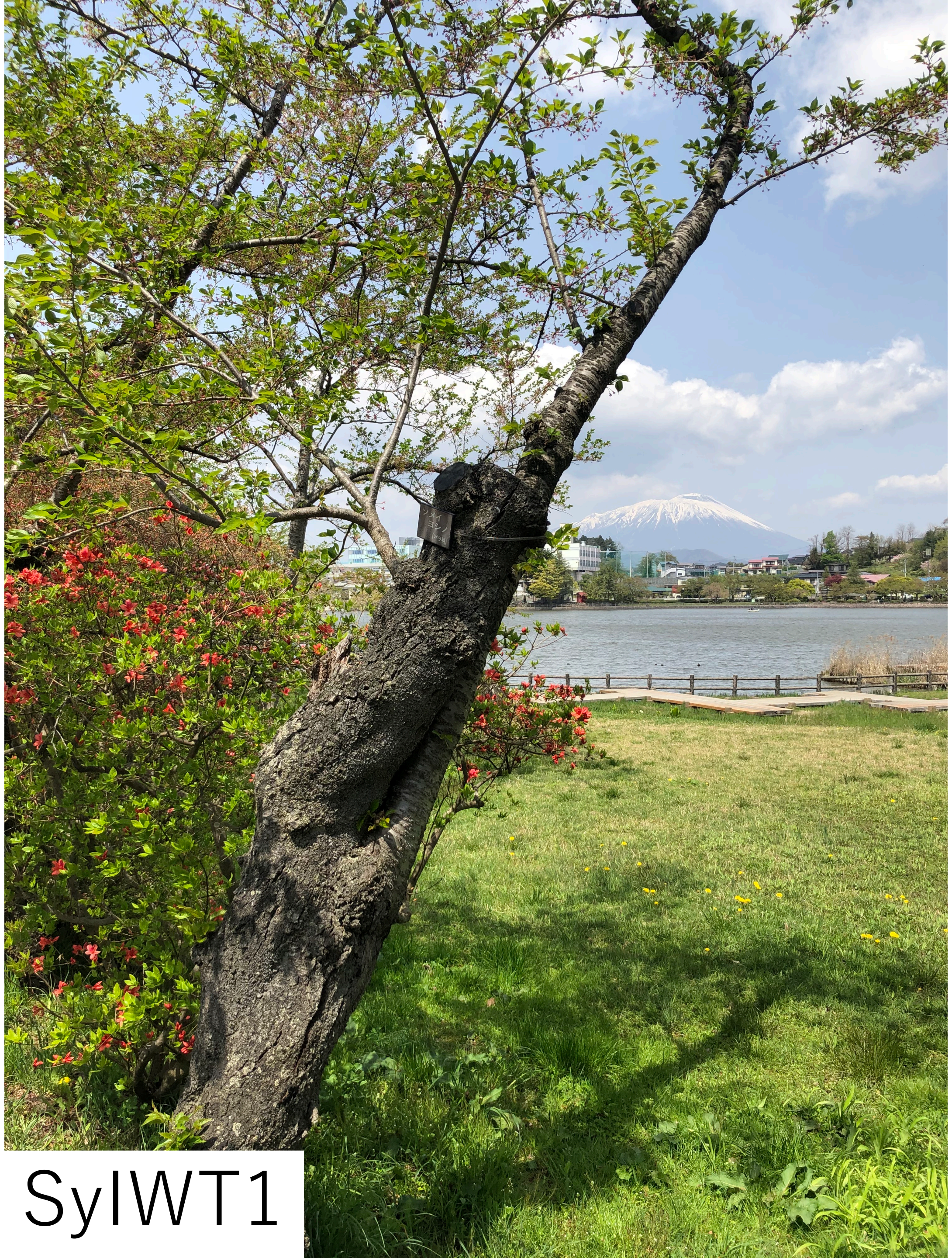
SyIWT1

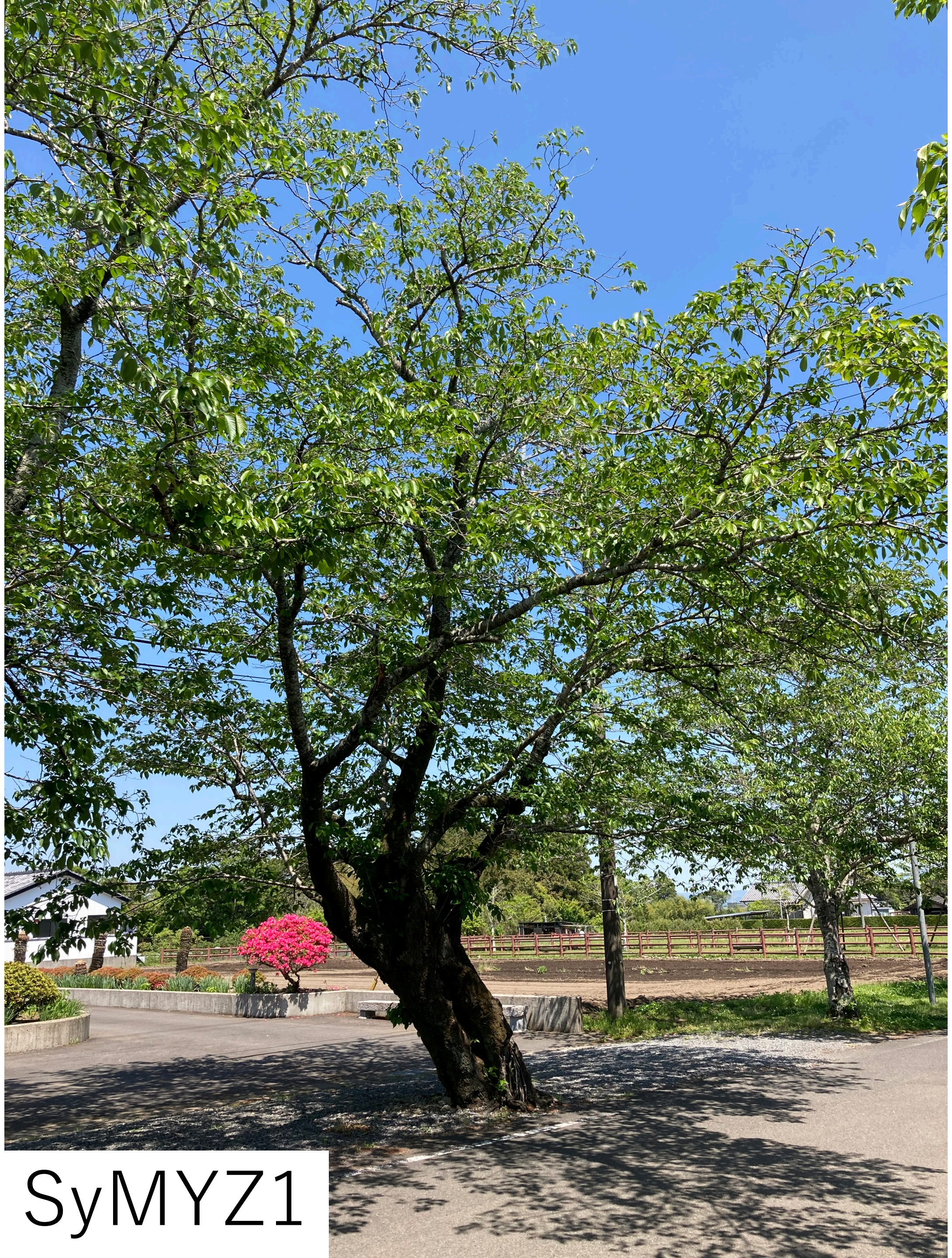

SyMYZ1

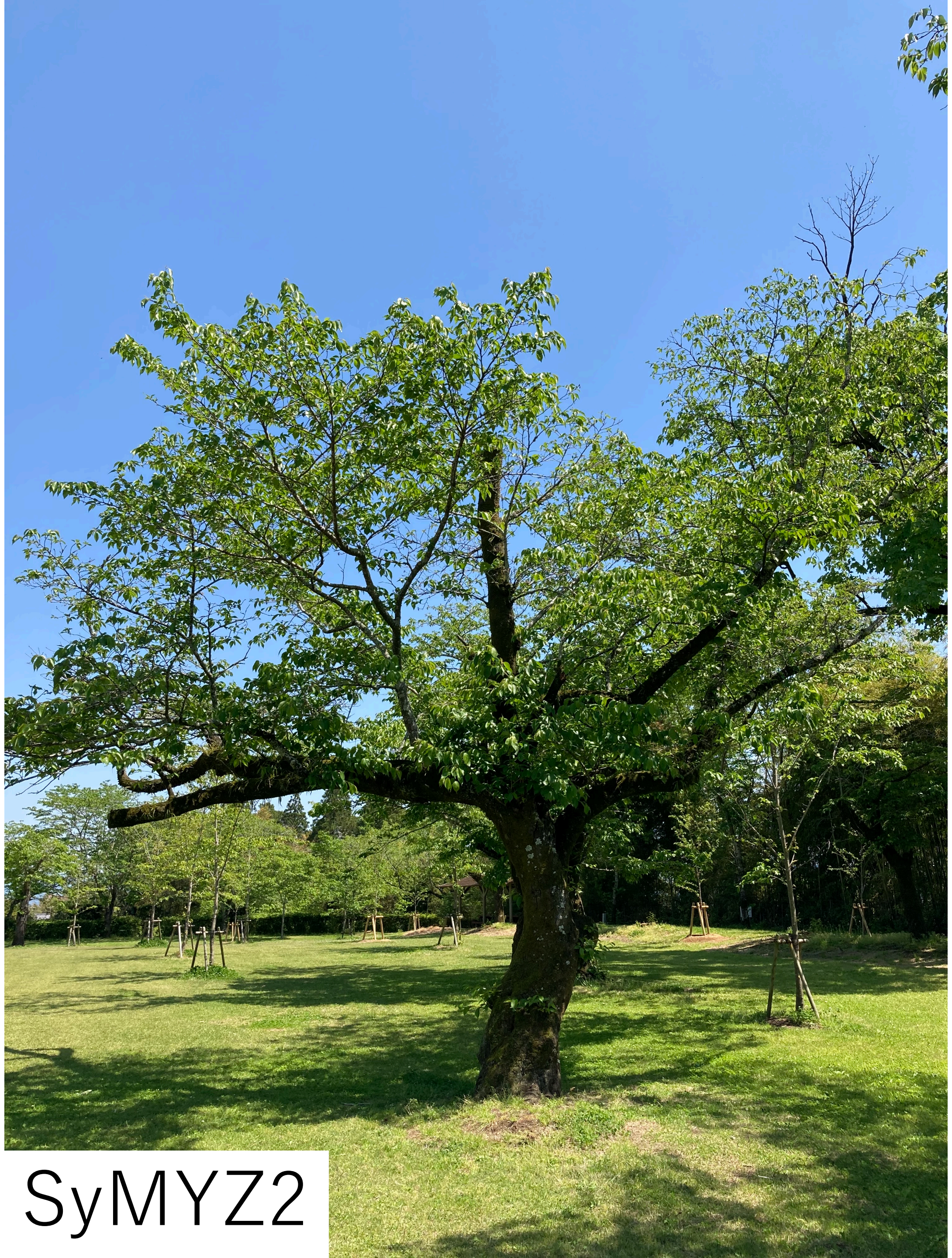

SyMYZ2

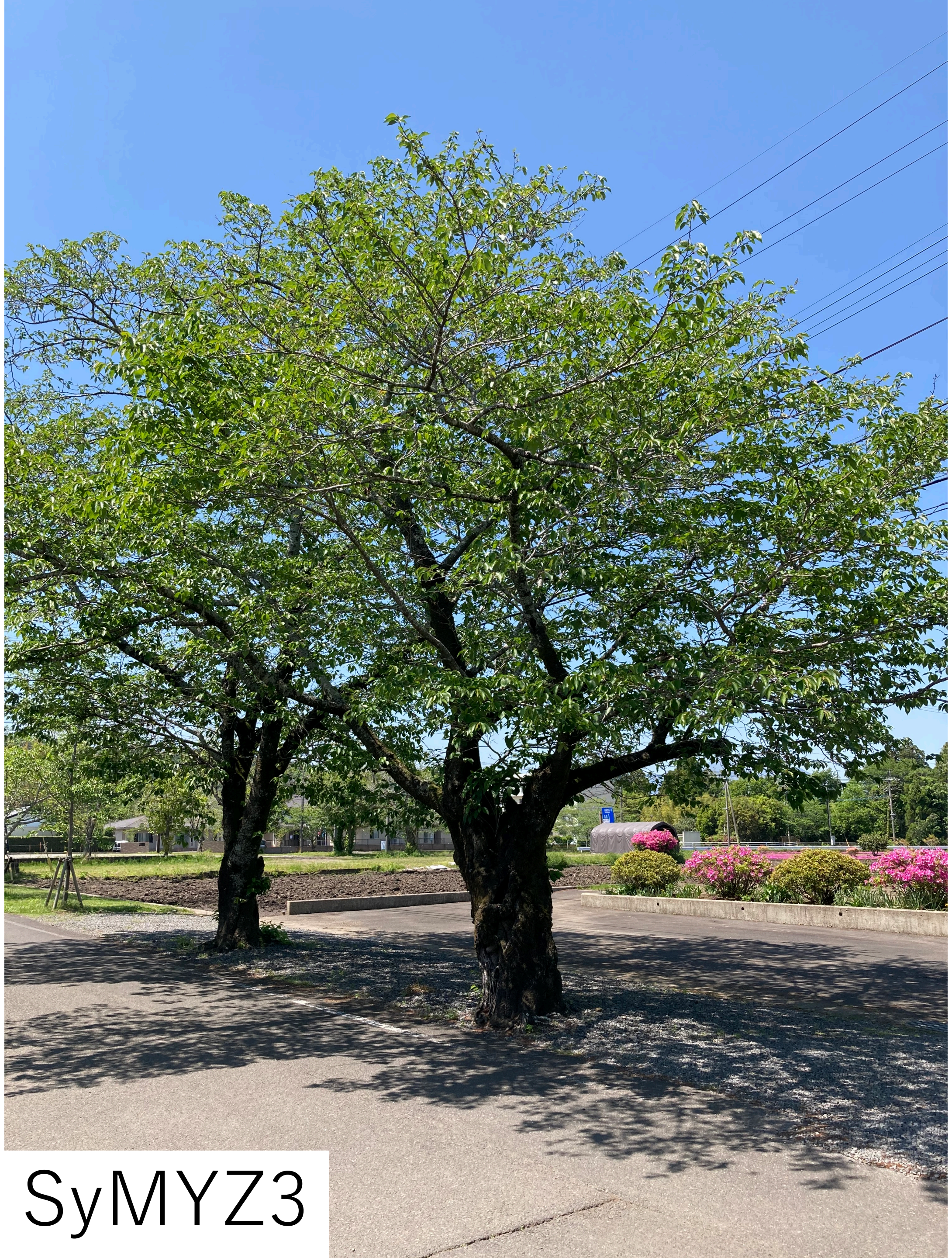

SyMYZ3

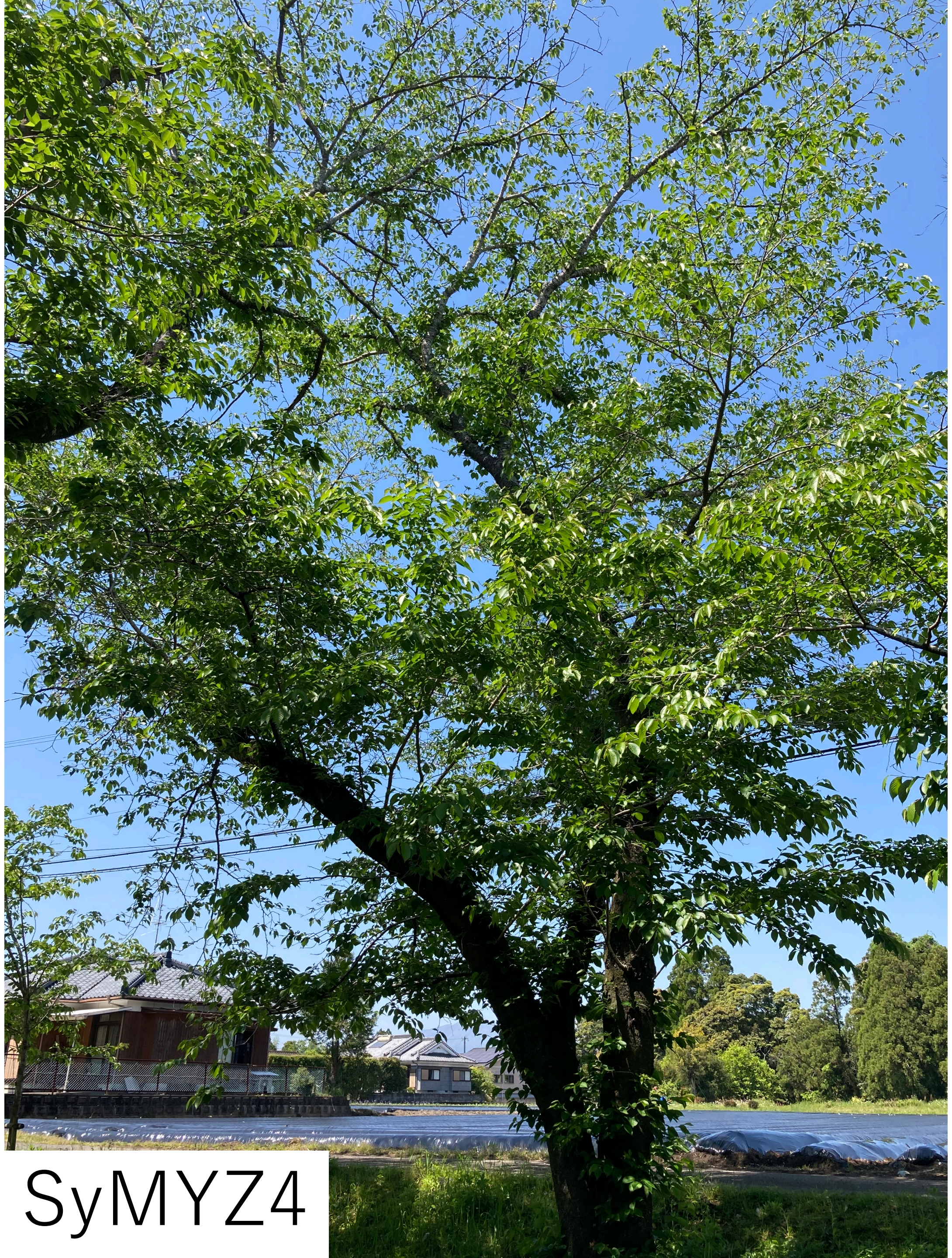

SyMYZ4

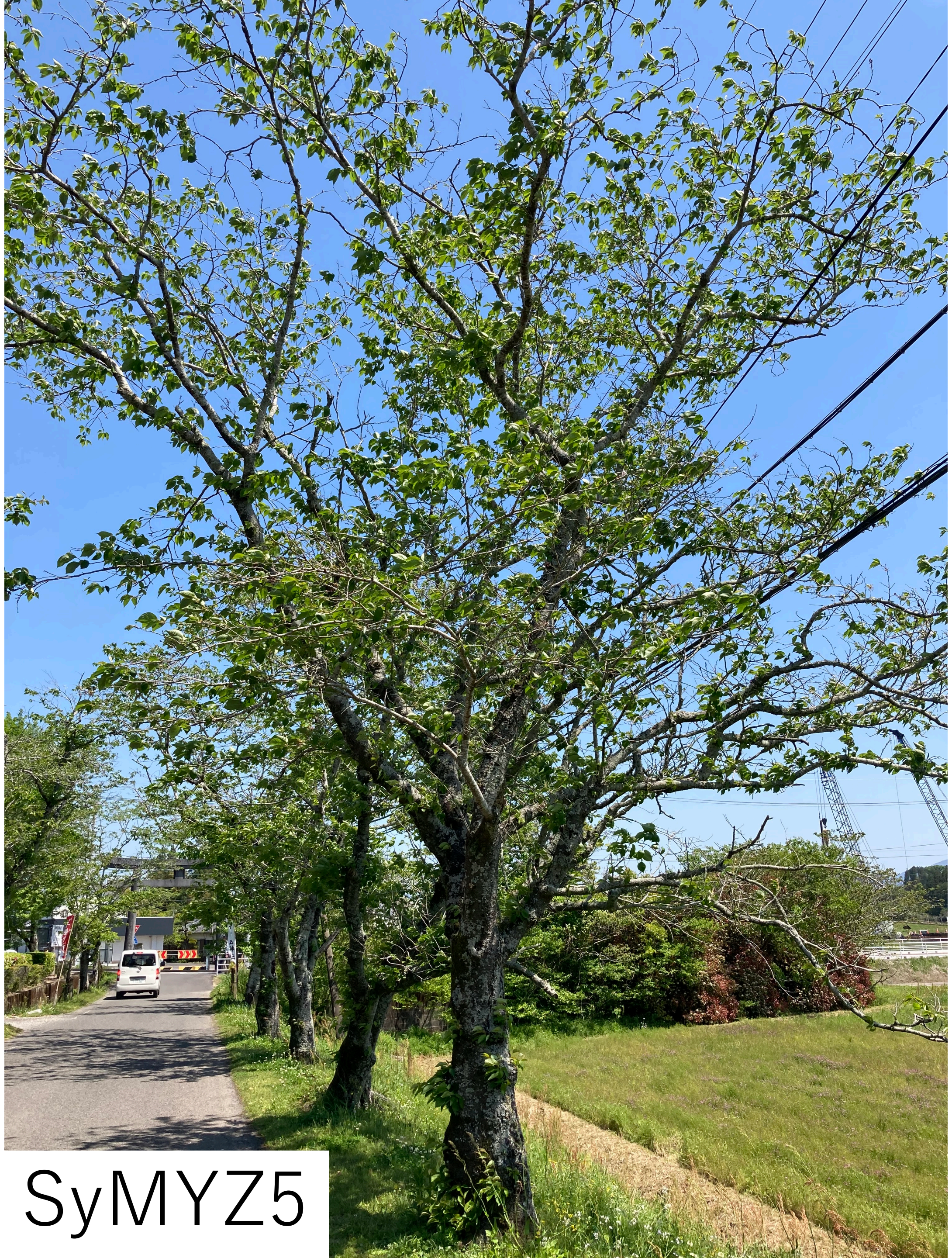

SyMYZ5

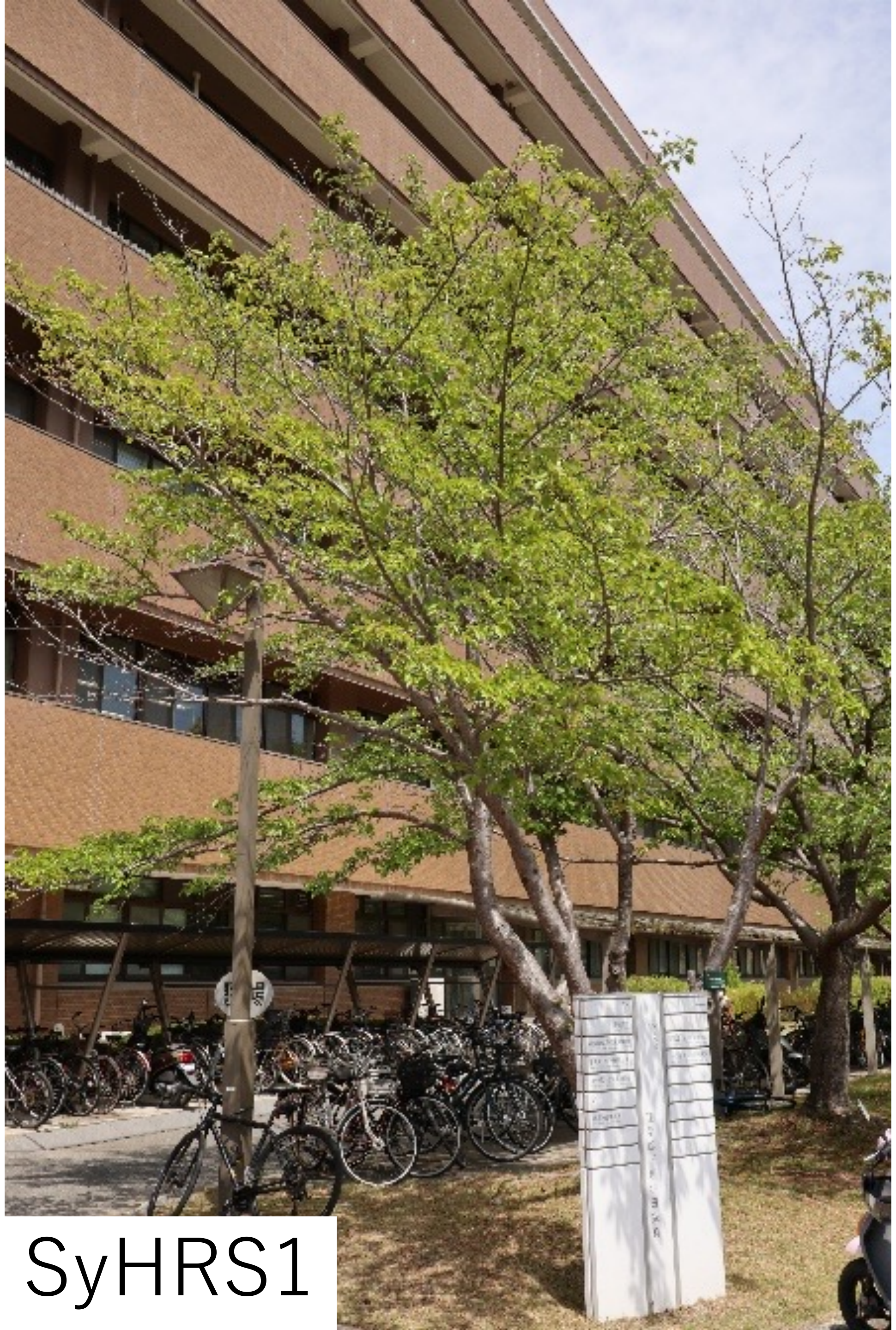

SyHRS1

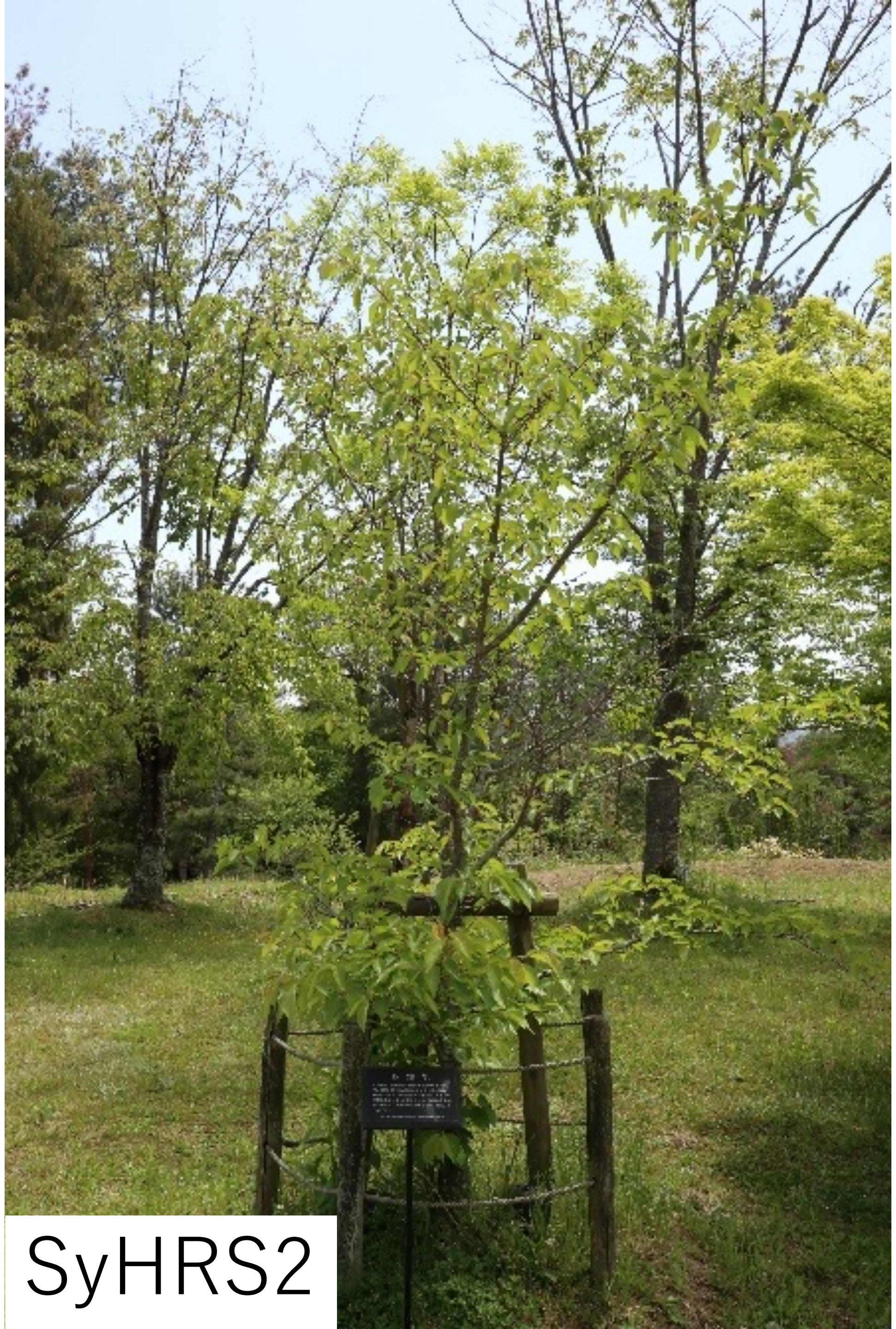

SyHRS2

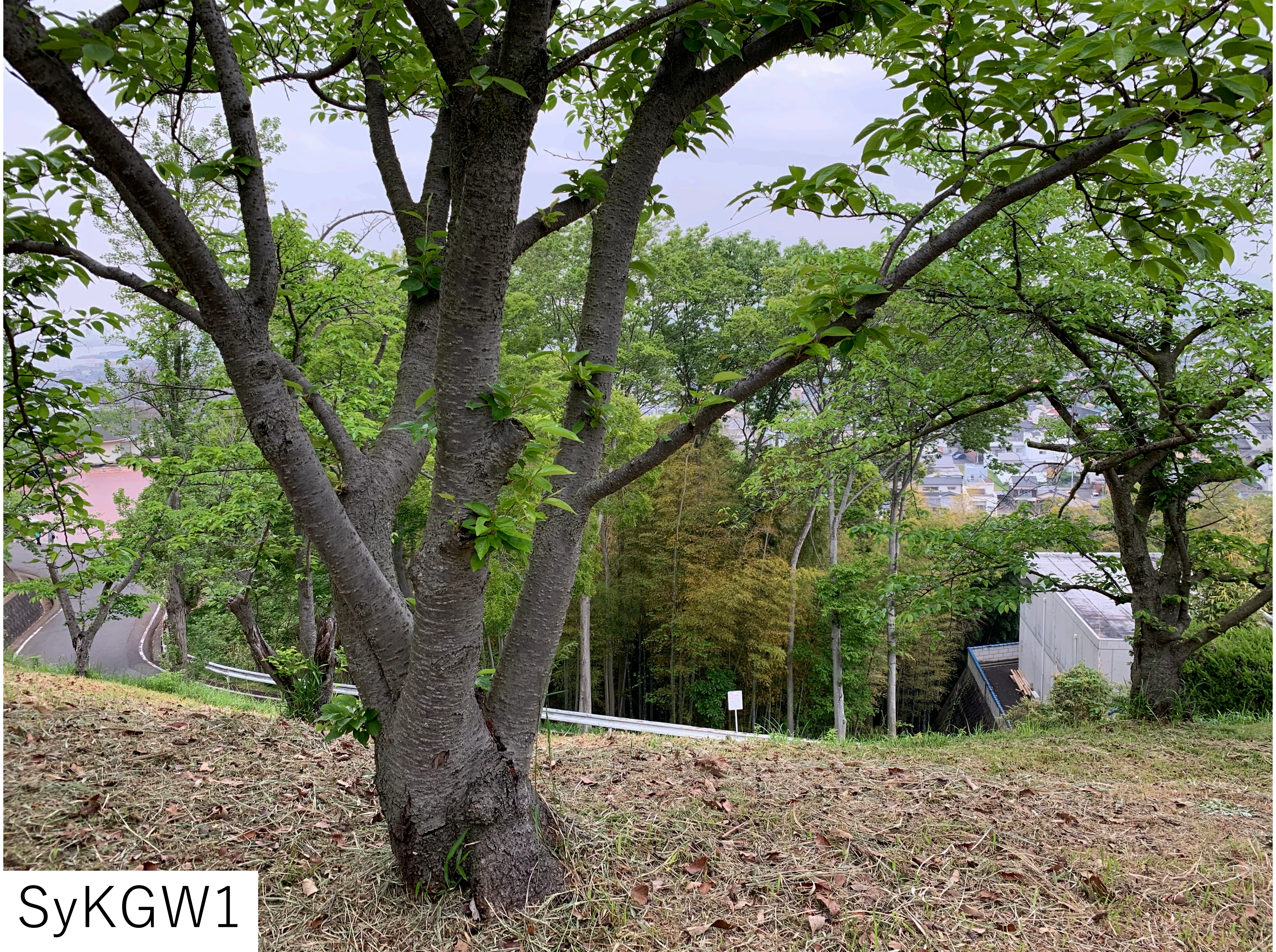

SyKGW1

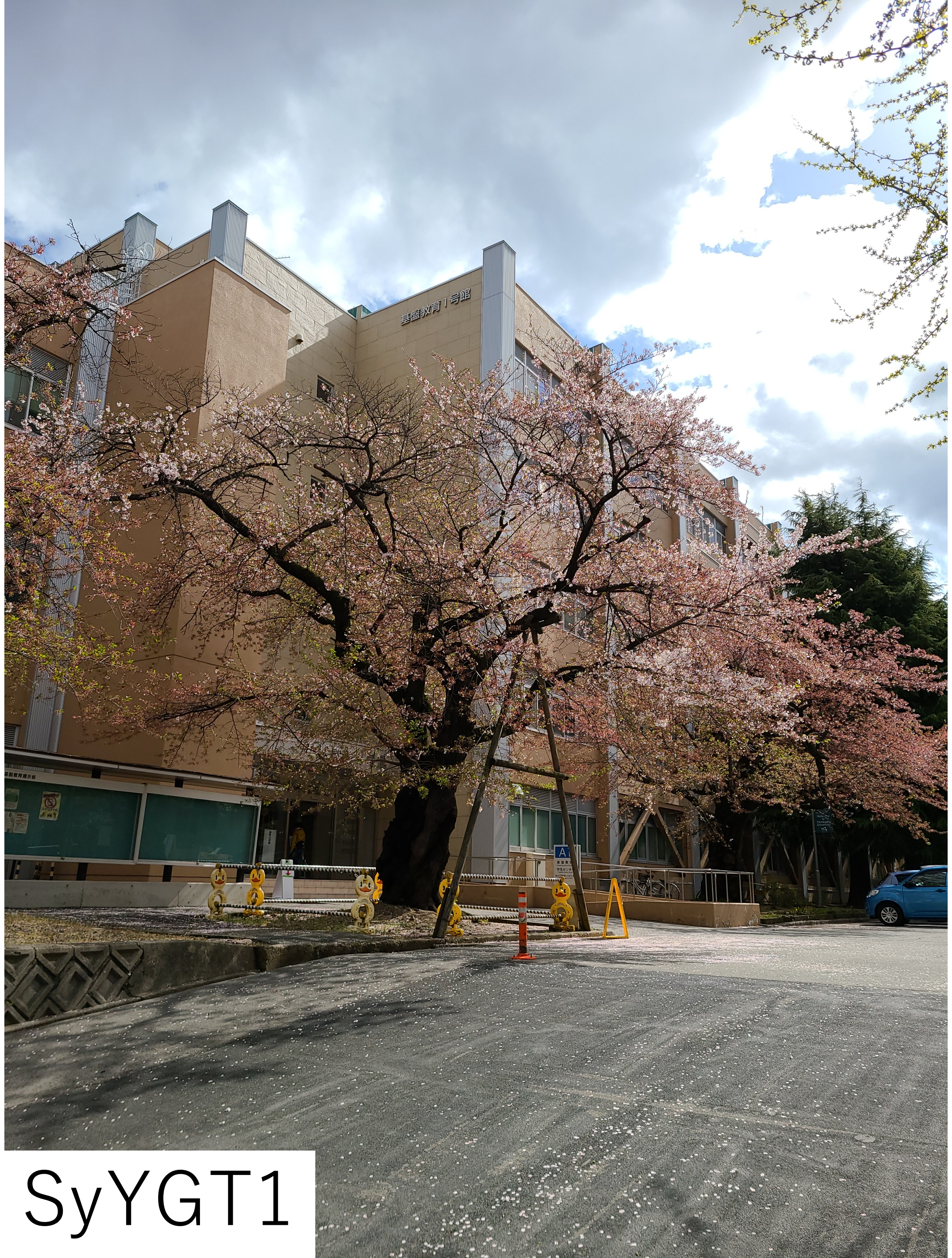

SyYGT1

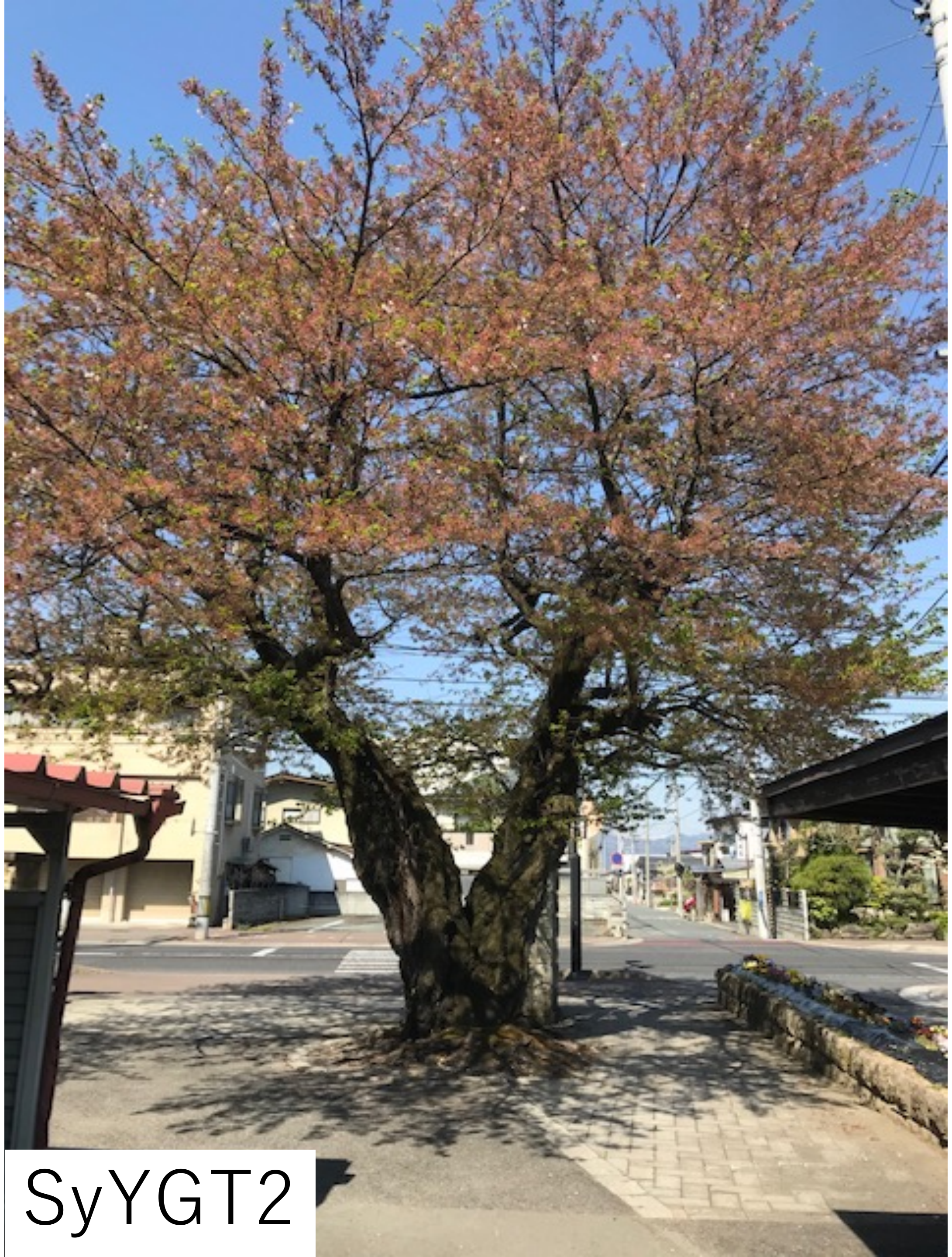

SyYGT2

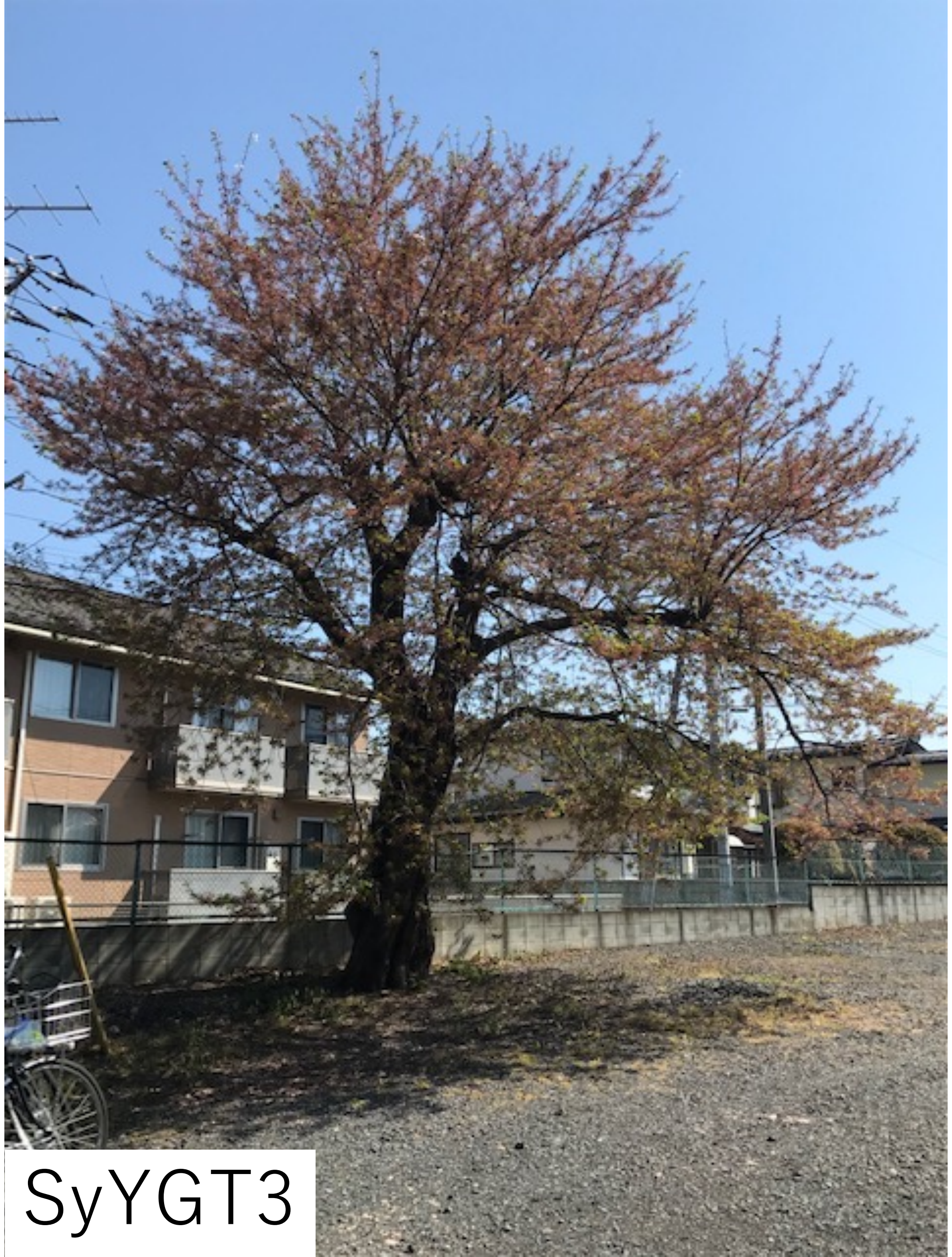

SyYGT3

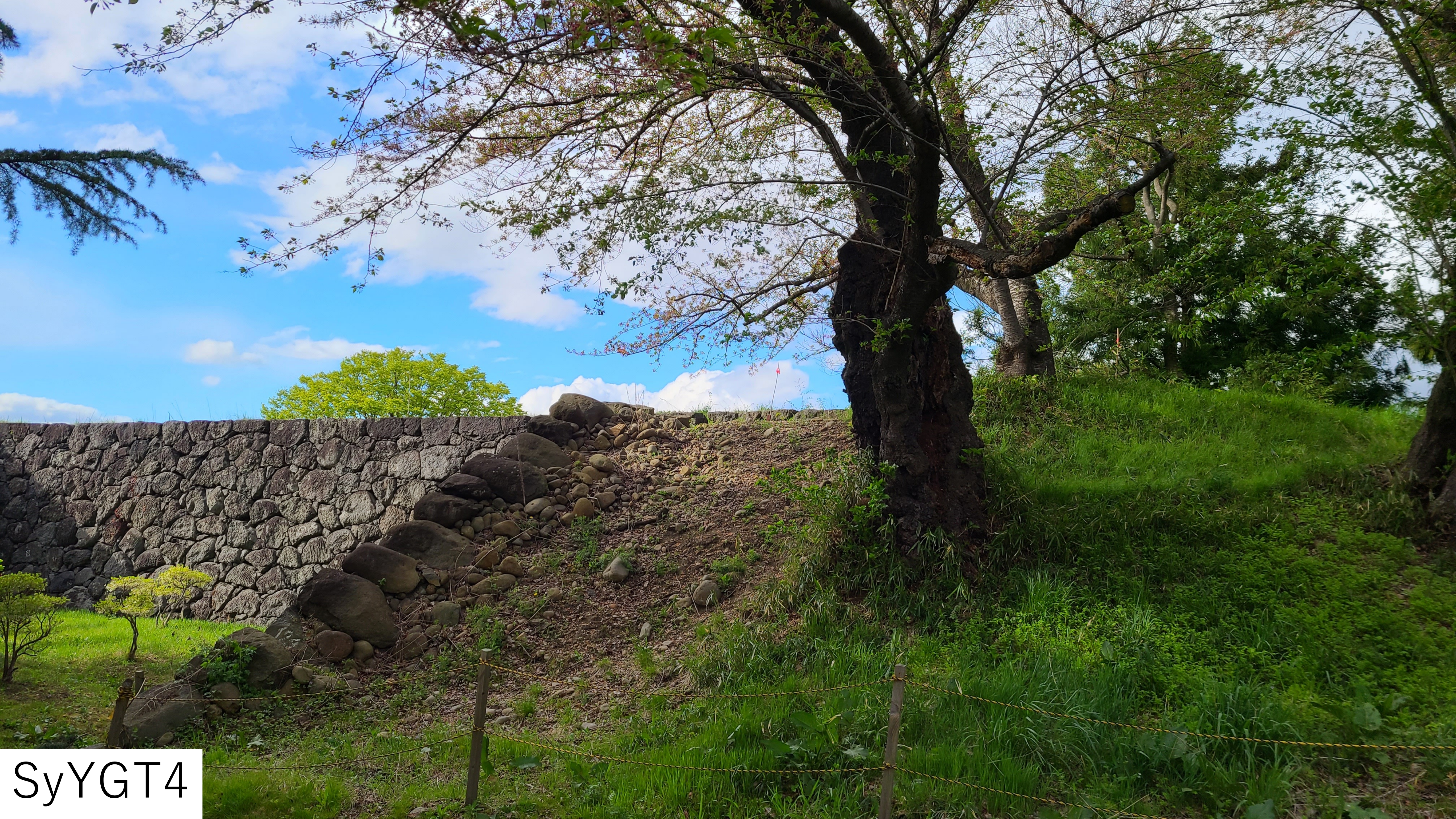

SyYGT4

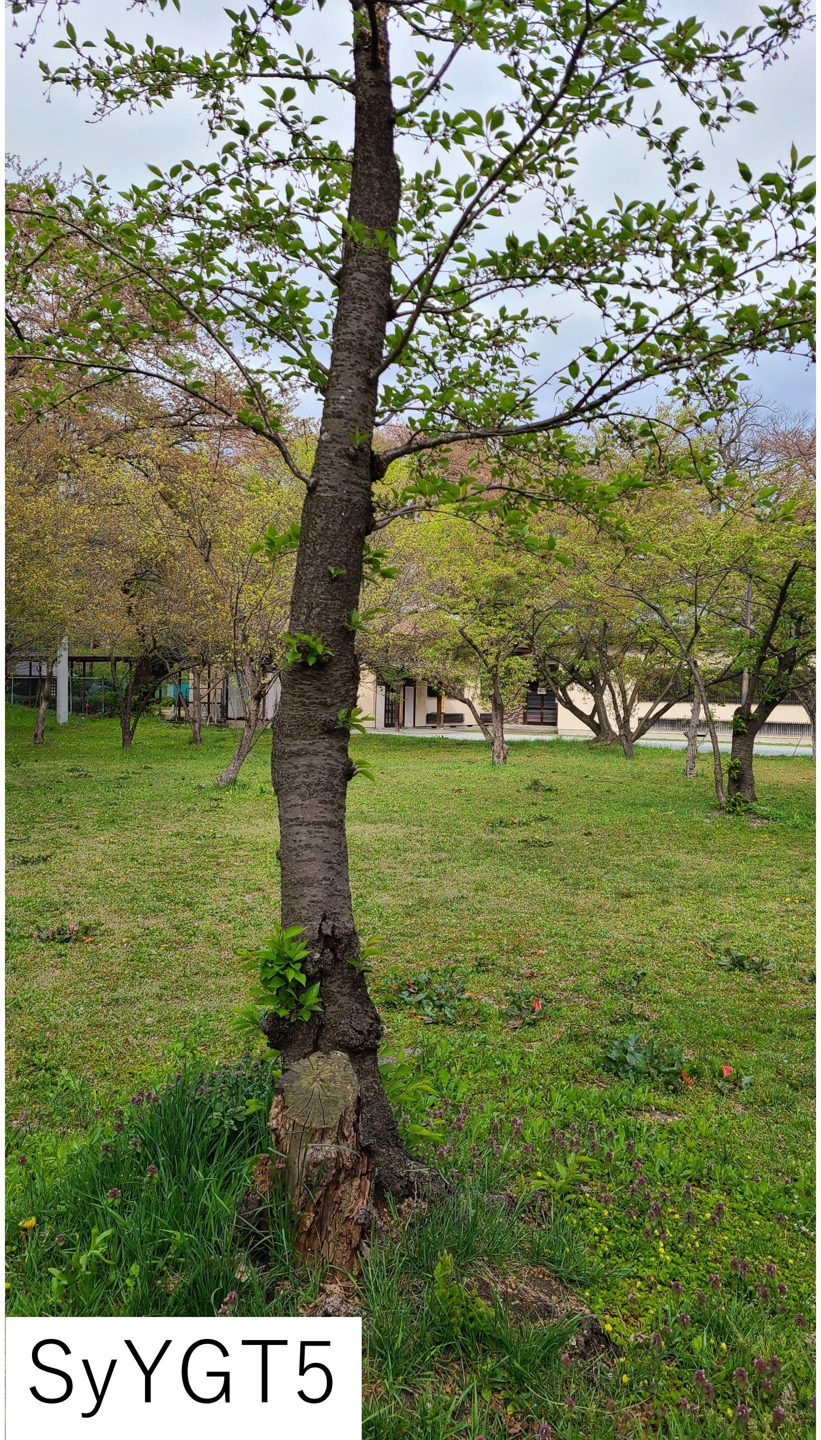
SyYGT5

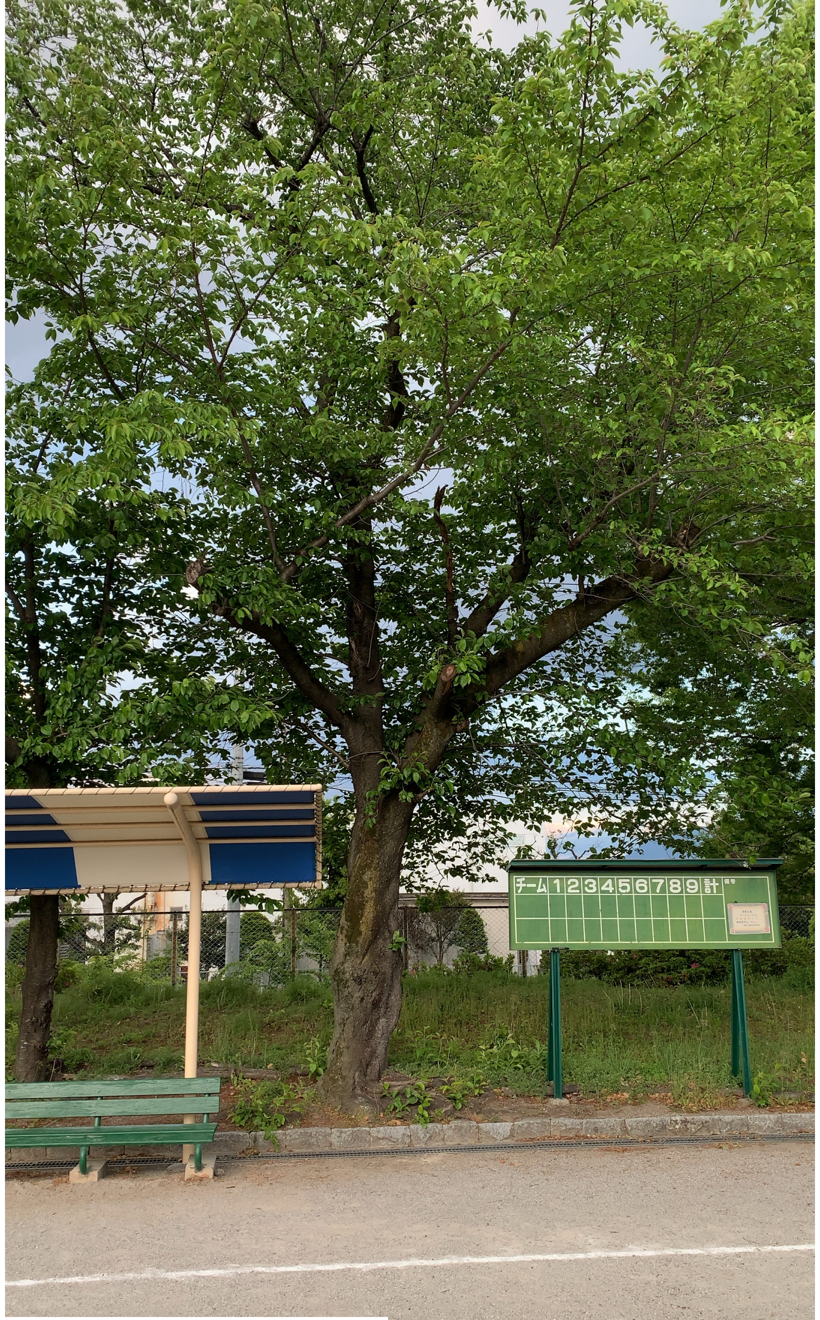


SyYMN1

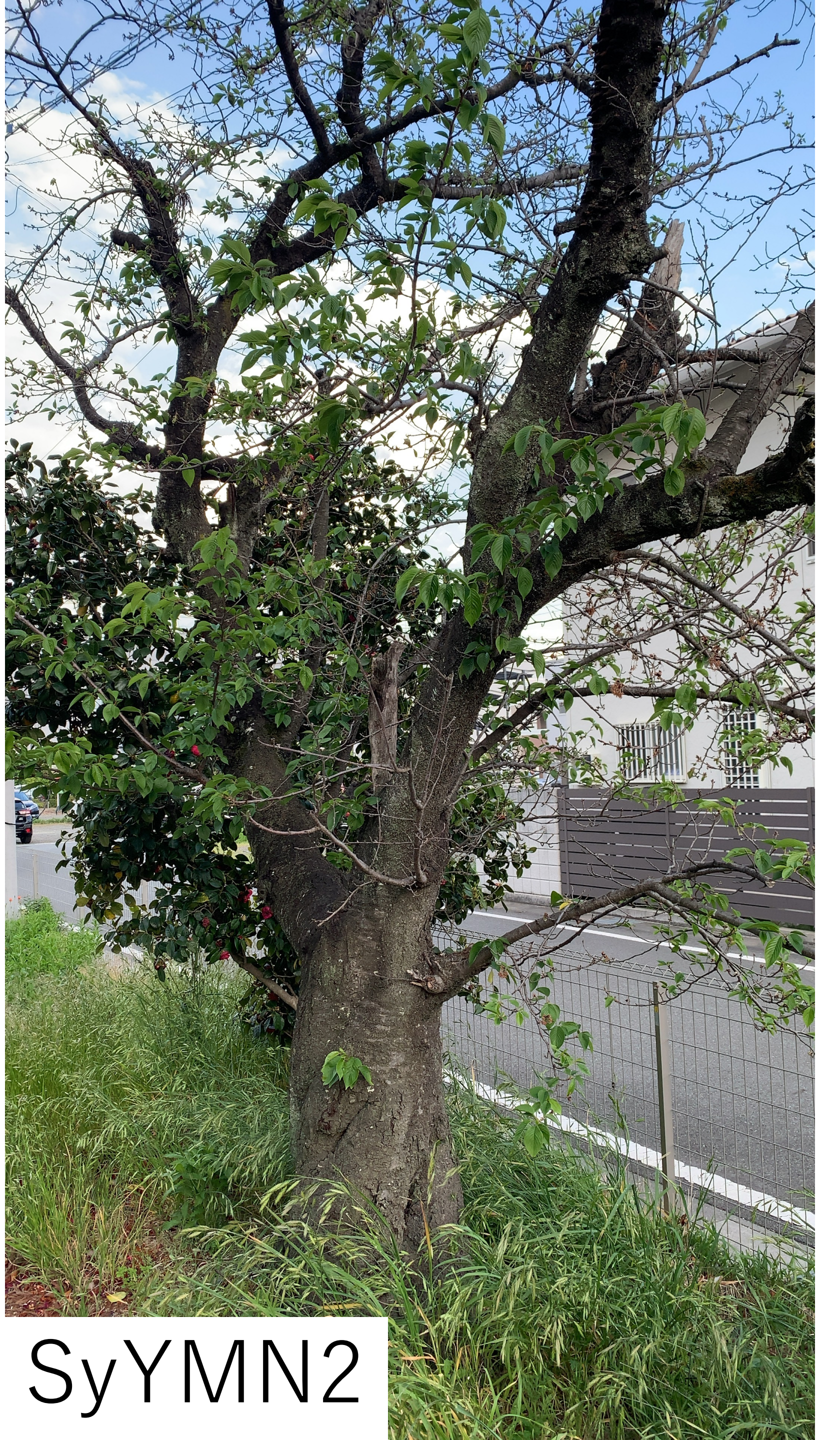

SyYMN2

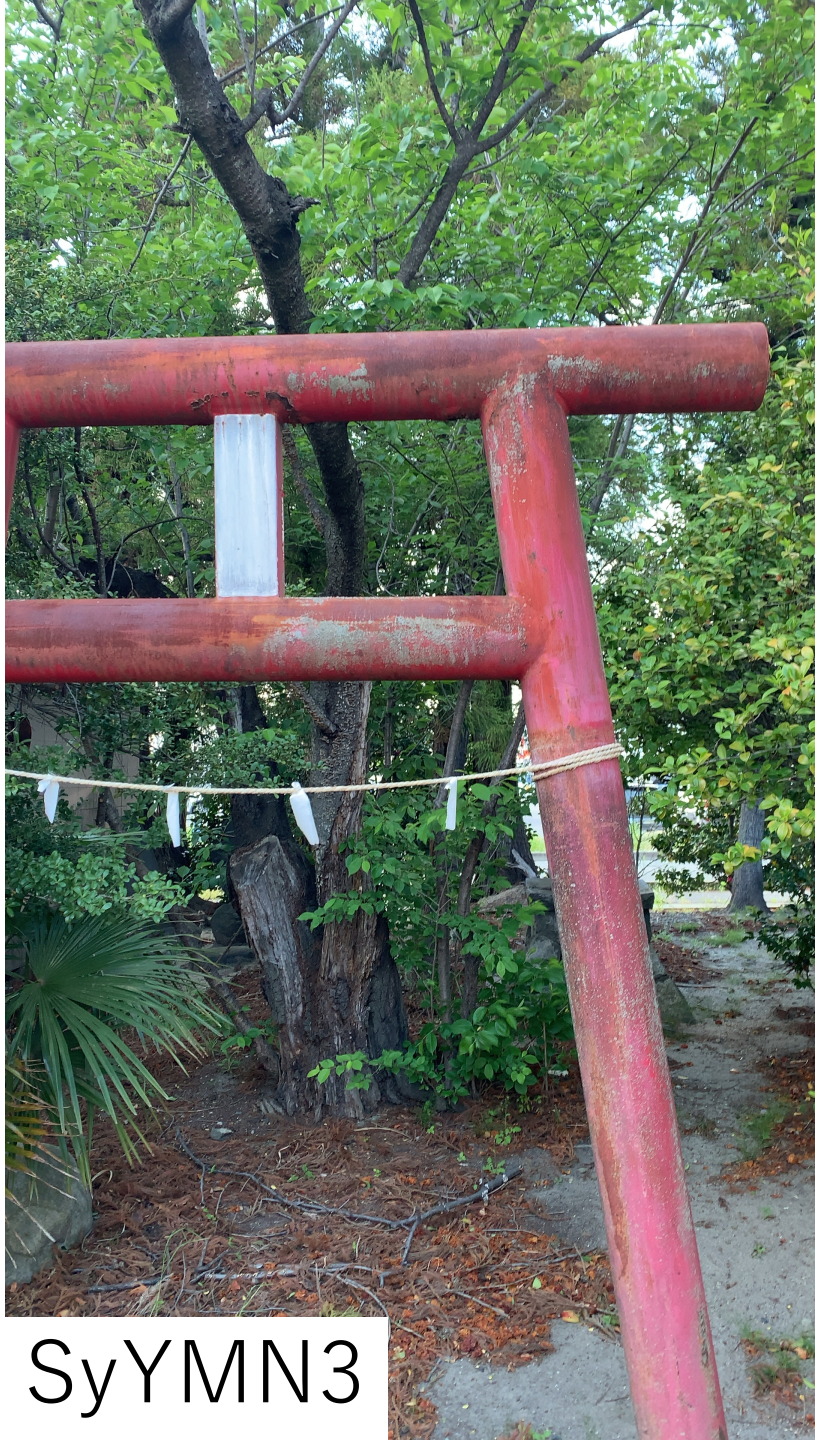

SyYMN3

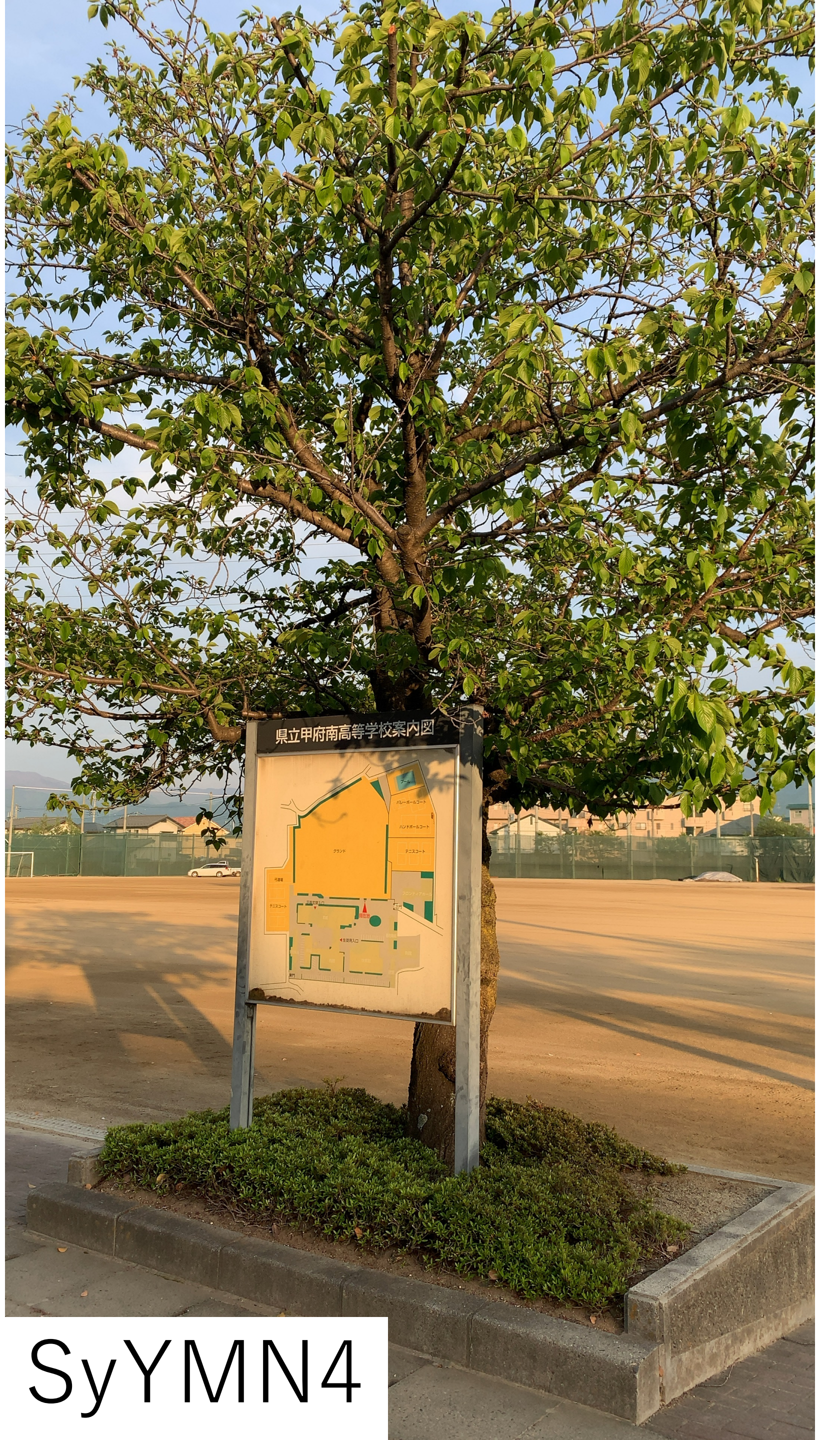


SyYMN4

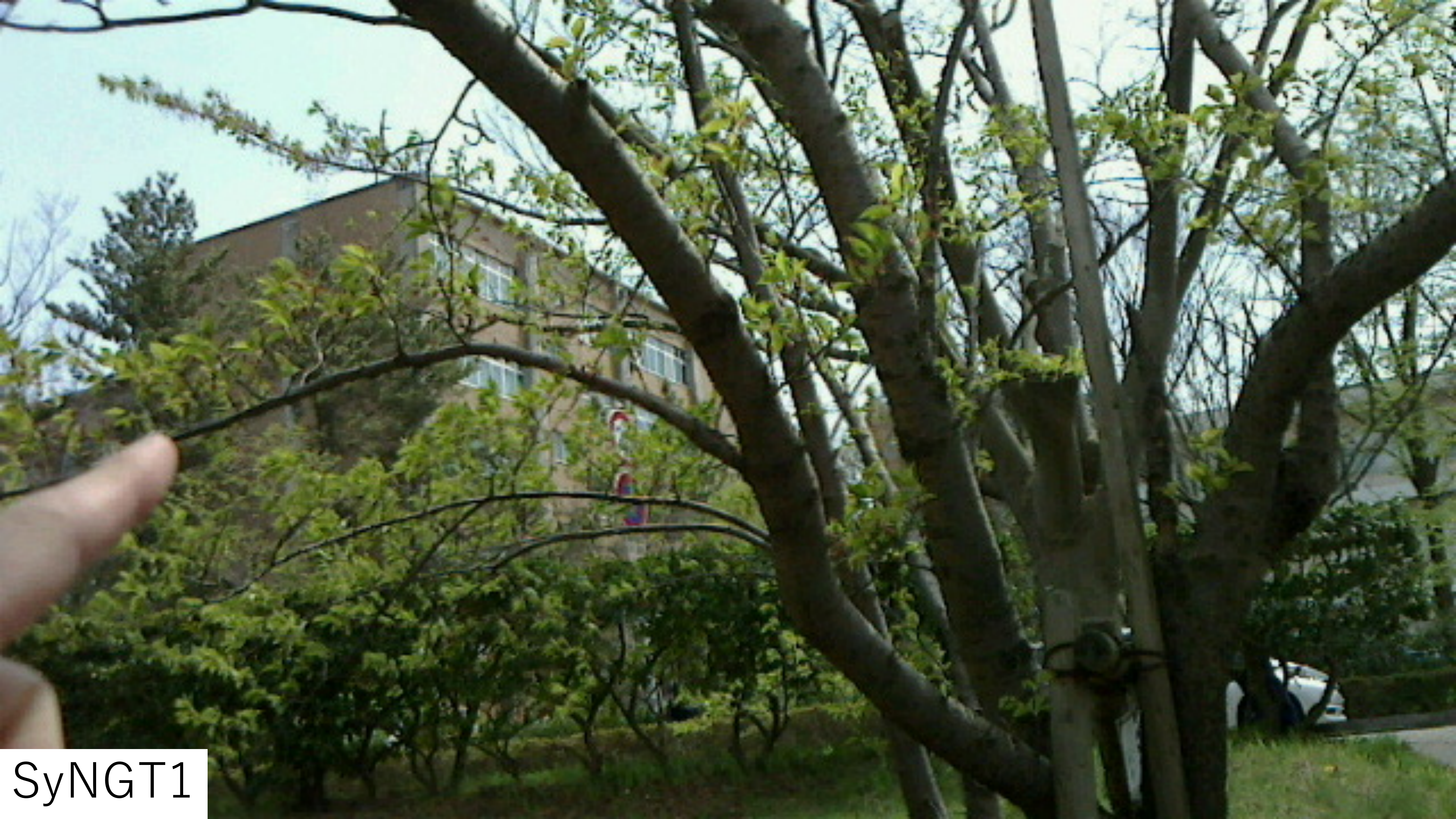
SyNGT1

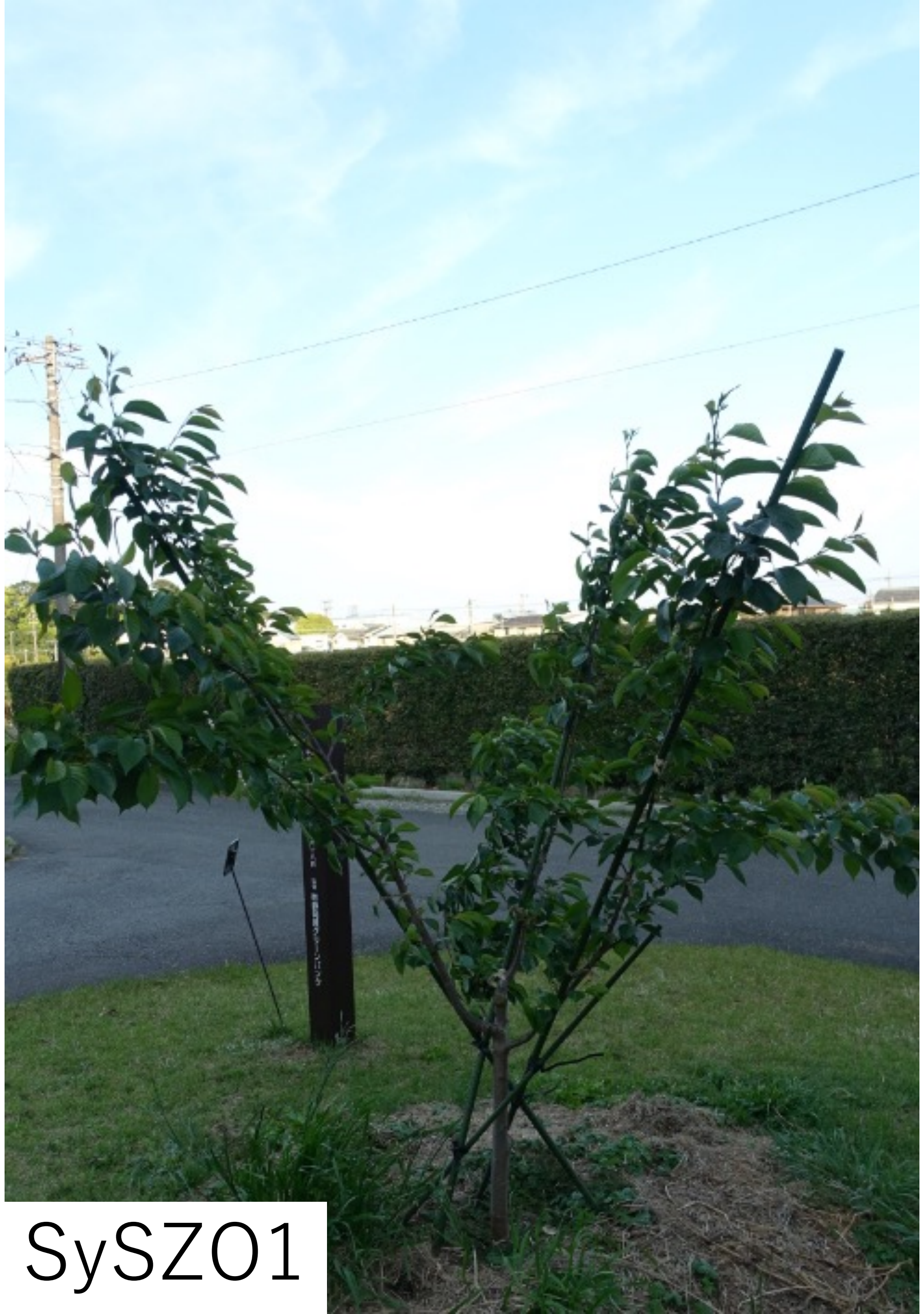

SySZ01

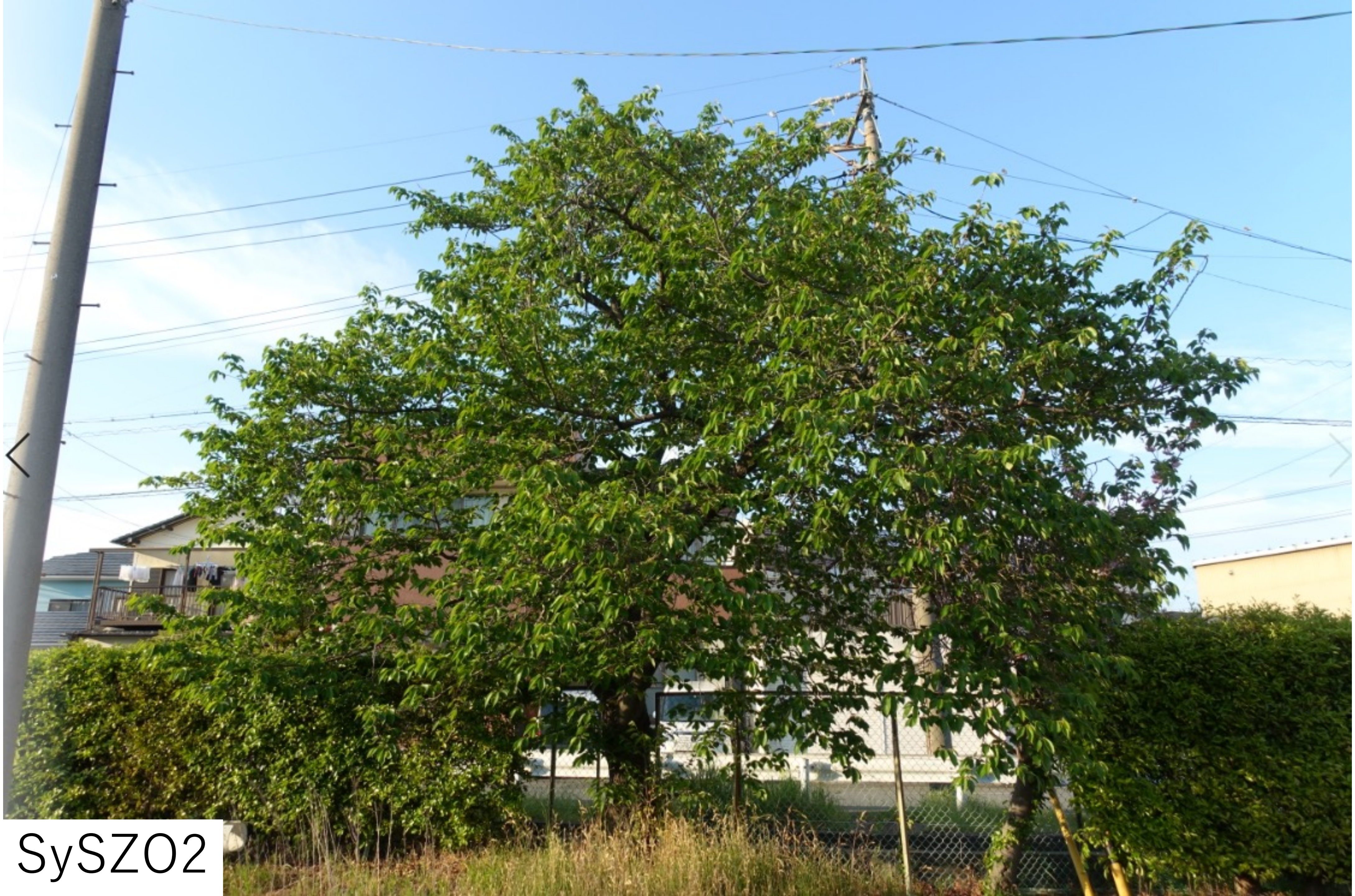

SySZ02

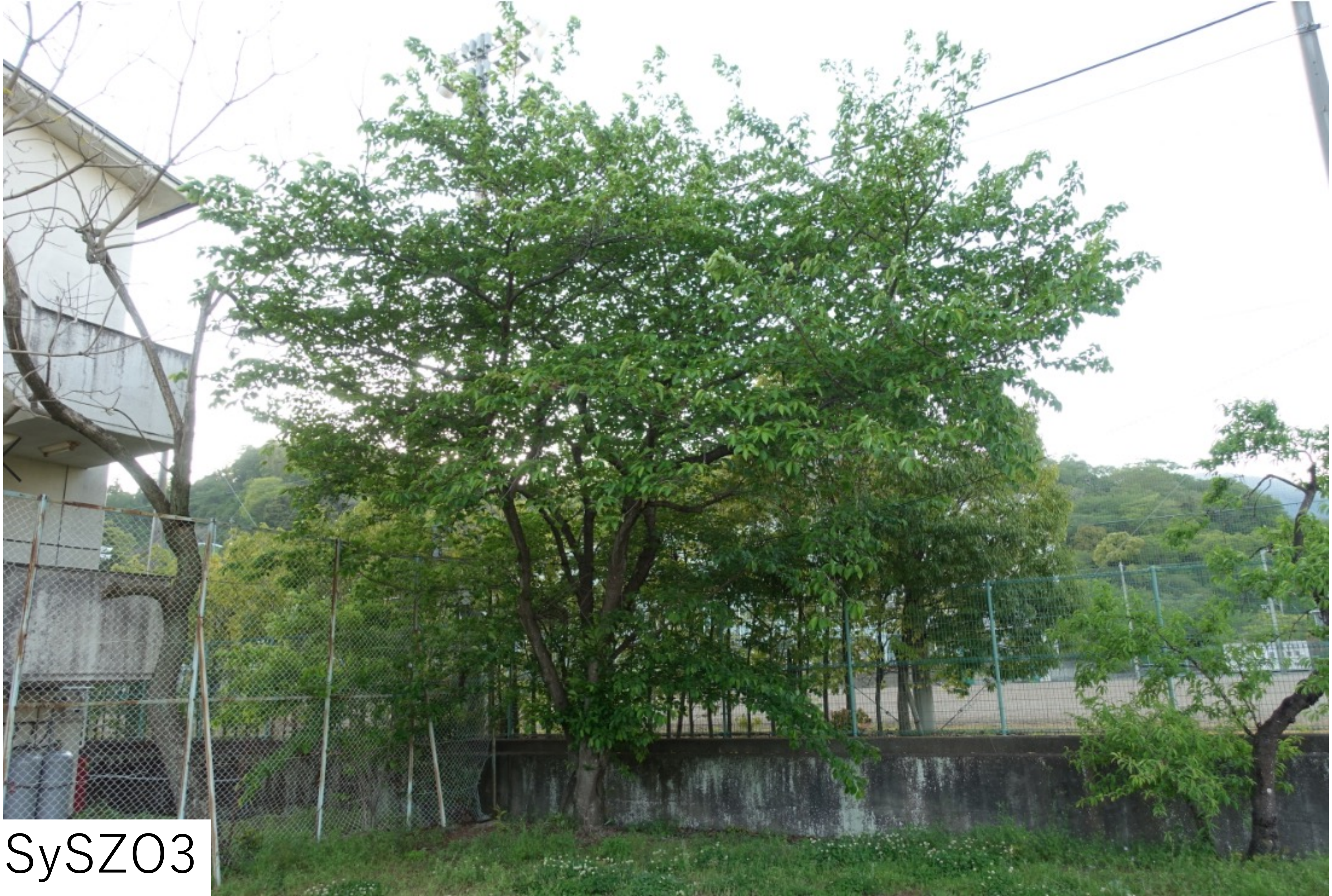

SySZ03

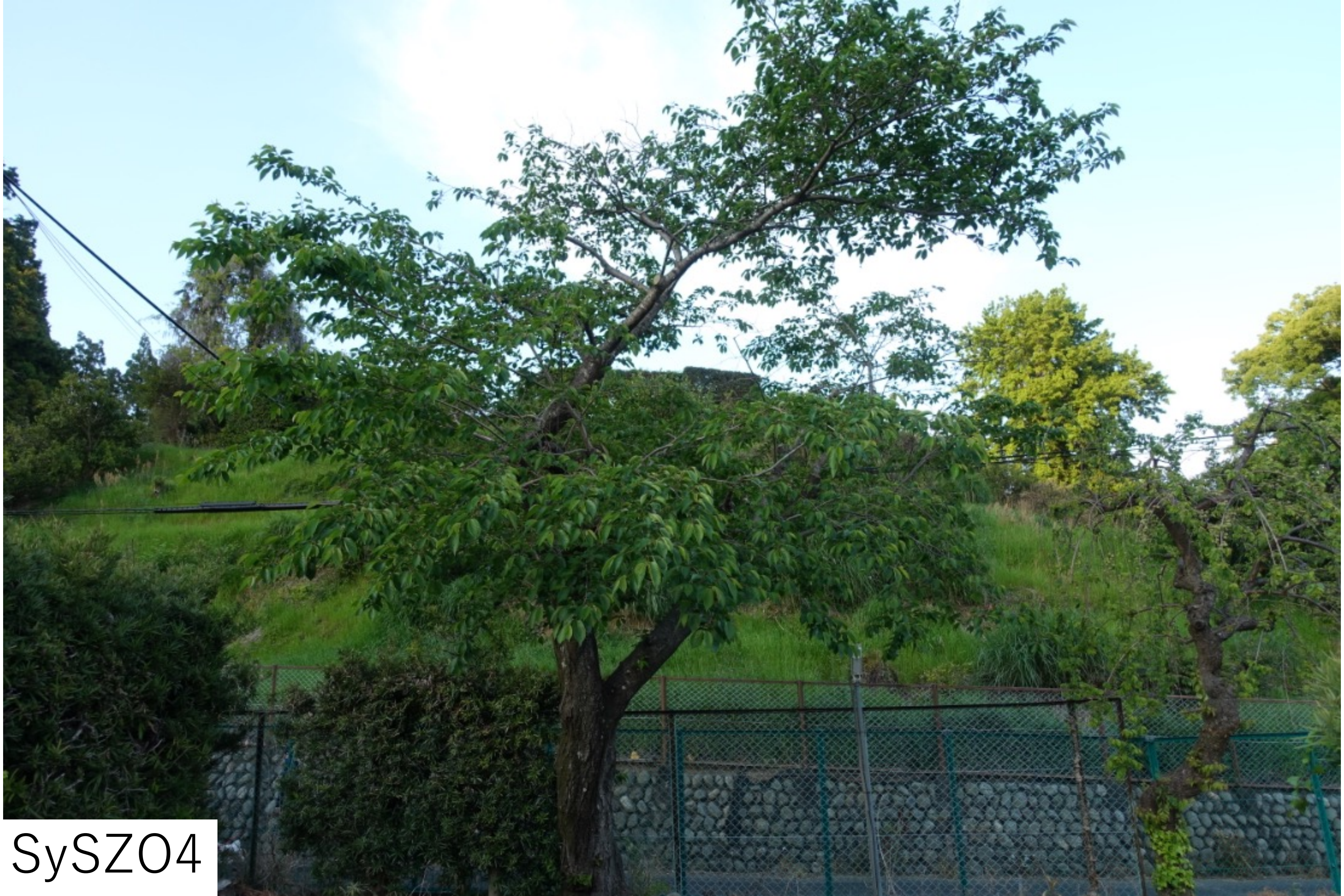

SySZ04

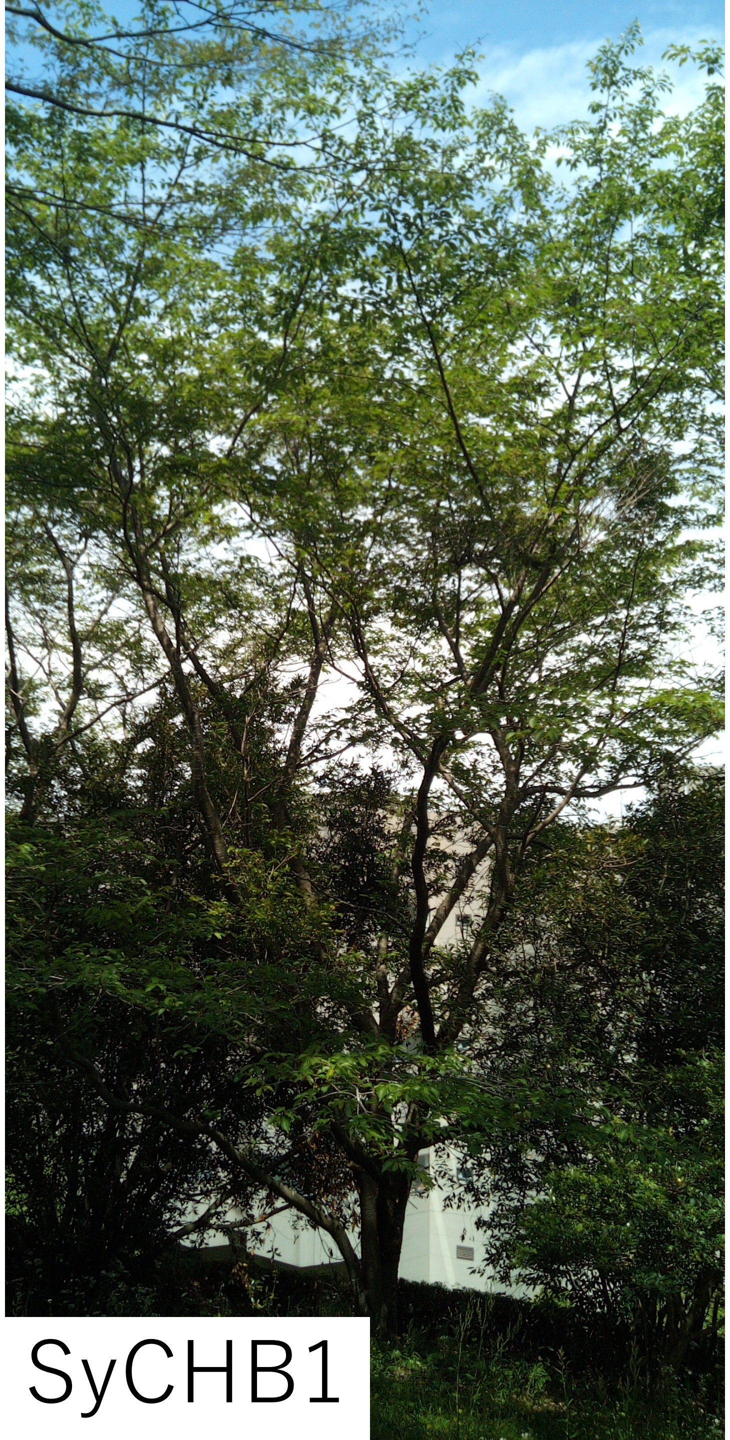

SyCHB1

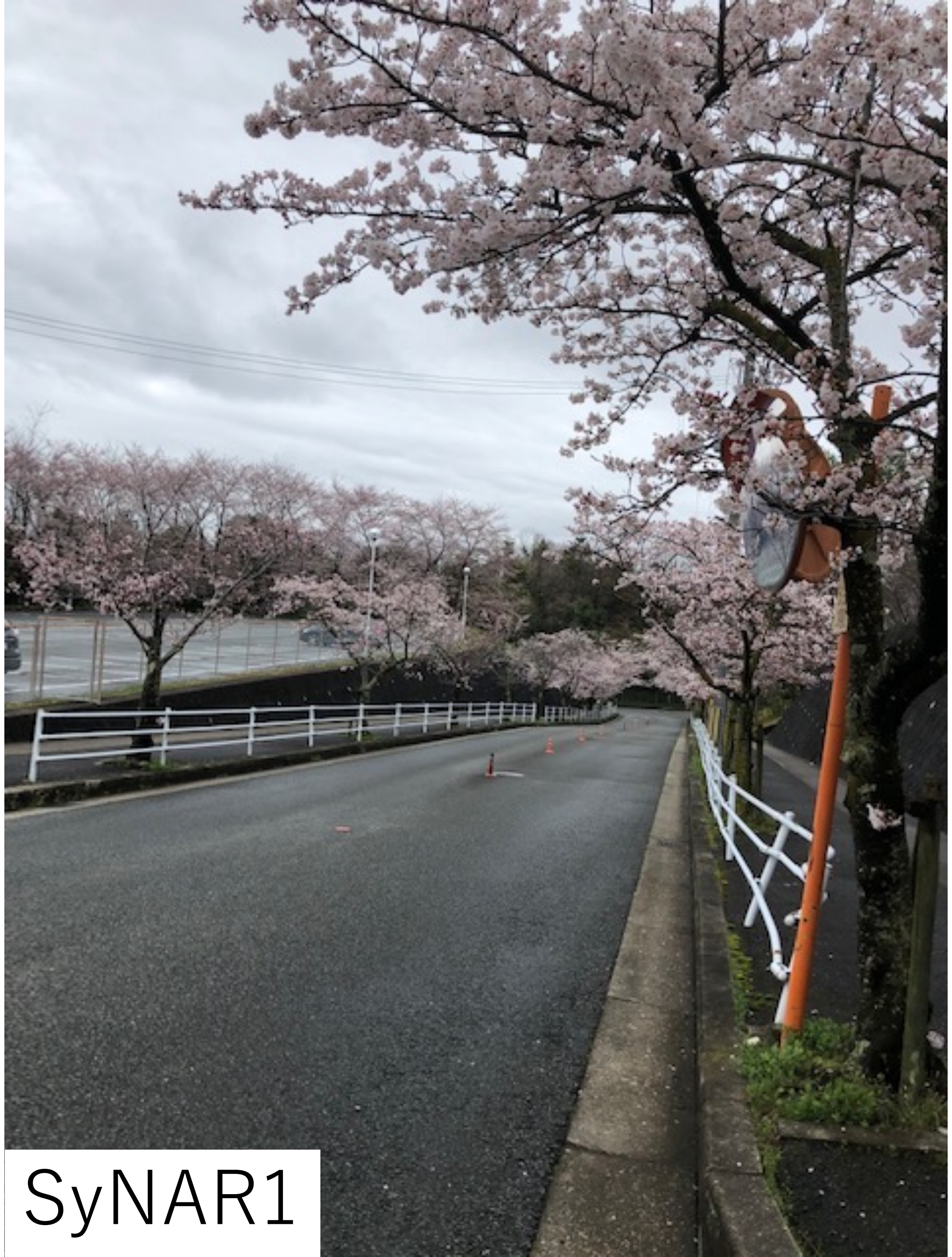

SyNAR1

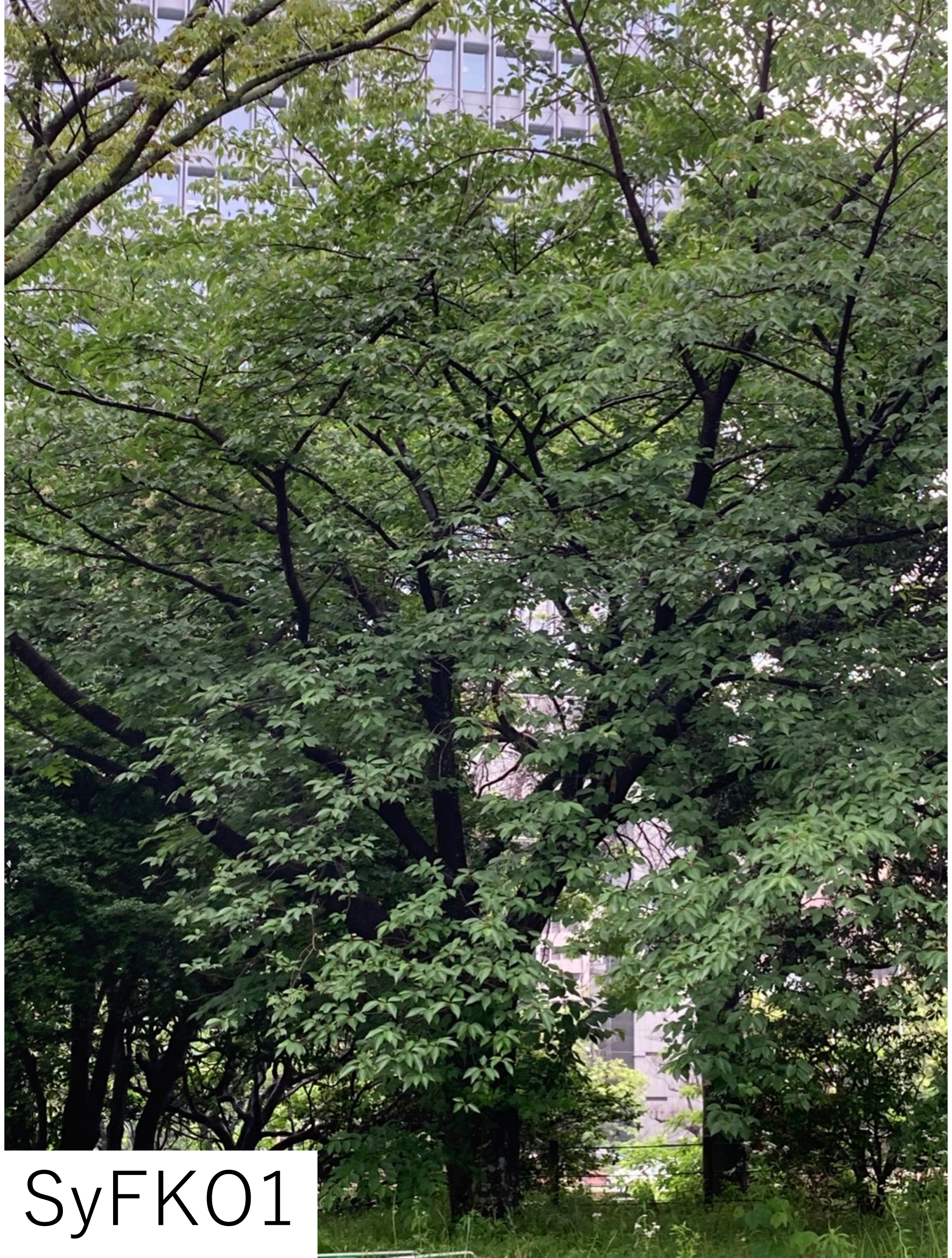

SyFK01
